# Effects of Left Atrial Wall Thickness on Myocardial Mechanics and Blood Dynamics using Multiscale Modeling

**DOI:** 10.64898/2026.06.09.731221

**Authors:** Boyang Gan, Lei Shi, Ian Y. Chen, Vijay Vedula

## Abstract

**Purpose:** Patient-specific models of left atrial (LA) mechanics often assume uniform left atrial wall thickness (LAWT), but the effect of LAWT on the mechanics and hemodynamics remains less quantified.

**Methods:** Four LA myocardium models were built from gated CTA images: a baseline variable thickness (VT#0), two reduced-dilation variants, and a 2mm uniform thickness model. Multi-scale mechanics and blood flow simulations were performed across all the thickness variants using model parameters personalized on the baseline model. Predicted displacements, wall stresses and strains, and hemodynamics were compared.

**Results:** Across all LAWT variants, myocardial volume spanned 14.4–19.9mL (38%), while cavity volume remained mostly within 5% of image data throughout the cardiac cycle. Circulatory system output, myocardial displacements, and strains varied by 5–6% relative to the baseline model. Instantaneous stresses increased by up to 19% in the thinner variable thickness models and decreased by up to 16% in the uniformly thick case. Globally, the area under low time-averaged wall shear stress (TAWSS) varied between 23% and 30% across all thickness variants, while LA exposed to elevated oscillatory shear index (OSI) increased from nearly 6% to 19%. Over 90% of LAA was exposed to low shear, but the high-OSI area increased from 7% in VT#0 to over 30% in Uniform.

**Conclusion:** A personalized multiscale modeling framework was leveraged to demonstrate that the left atrial myocardial stresses and oscillatory shear had a greater sensitivity to local wall thickness representation compared to cavity volumes, tissue displacements, strains, and mean blood shear.

## 1 Introduction

The left atrium (LA) plays a central role in left ventricular filling and acts sequentially as a reservoir, conduit, and booster pump over each cardiac cycle [1–3]. It also has a complex anatomy and tissue structure, including asymmetric pulmonary venous inflow, a compliant and trabeculated left atrial appendage (LAA), and a marked heterogeneity in wall structure [4–6]. LA dysfunction is closely linked to atrial fibrillation (AF), the most prevalent arrhythmia affecting over 50 million people worldwide [7], and a major contributor to stroke, heart failure, and cardiovascular mortality [8, 9]. Arising from coupled electrical, structural, and hemodynamic remodeling, AF is also strongly associated with blood stasis and thrombus formation, particularly in the LAA cavity, increasing the risk of stroke by 4 to 5 fold [10–13]. Because blood stasis is governed in part by the interaction between atrial deformation and intracavitary blood transport, LA structure and wall motion are important determinants of the hemodynamic conditions leading to thrombus formation and the associated clinical risk [6, 14–17].

Recent advances toward cardiac digital twins are enabling the development of personalized physics-based and data-driven computational models tailored to an individual’s anatomy and physiology, including loading conditions and tissue properties [18, 19]. Such models provide a systematic framework for integrating multimodal clinical data, testing new mechanistic hypotheses, and evaluating biomechanical and hemodynamic responses to interventions [20]. In the left atrium, we developed a method to personalize myocardial passive mechanics using inverse finite element analysis (iFEA) [21]. The method has been extended to personalize left atrial mechanics and the circulatory system, aligning with clinical data, using a multistage personalization framework [22]. The left atrial mechanical digital twin provides estimates of regional tissue stress and strain, as well as subject-specific wall motion, which is then used to drive blood flow [22]. The resulting hemodynamics, including markers of blood stasis such as residence time and shear forces, could be used to assess thrombogenic risk and treatment efficacy [15, 23–33].

However, one critical modeling input that remains poorly constrained is the spatial distribution of the left atrial wall thickness (LAWT). Most existing electromechanical and fluid-structure interaction models of the left atrium assume a uniform myocardial thickness [34–36], even though physiological thickness varies spatially from 1 to 4mm [5, 37]. This simplification is largely due to the difficulty of delineating the epicardial boundary on medical images and, as such, the thickness extraction remains technically challenging [37–41]. At the same time, the uniform thickness assumption contradicts the known LAWT heterogeneity, demonstrated across multiple imaging and anatomical studies [37, 39, 42–44].

Protocols have been developed and tested across multiple patients to segment thick LA myocardium from contrast-enhanced computed tomography angiography (CTA) images [40]. Laplace solution-based methods were then developed to create a map of LAWT over the entire surface [40]. These are supplemented with benchmarking efforts for atrial wall segmentation and thickness measurement [42]. However, small segmentation differences at the voxel level can alter LAWT estimates, as well as chamber volume and surface curvature – key geometric features that will likely affect tissue mechanical response and blood flow patterns. These discrepancies between observations and model assumptions lead to a basic question: how sensitive are left atrial tissue mechanics and hemodynamics to wall-thickness assumptions and segmentation uncertainty? The current work aims to address this question using multiscale modeling.

A few computational studies have explicitly assessed the effects of wall thickness on LA biomechanics and function. Simulations of left atrial myocardial mechanics have shown that the wall stress depends on geometry and loading, and that patient-specific wall thickness and curvature can modulate regional stress distributions [34, 45–47]. In parallel, image-based computational investigations have demonstrated that the LA wall motion, appendage morphology, fibrosis, and patient variability can influence intra-atrial blood transport and hemodynamic surrogates of thrombogenicity [15, 23–32, 48–50]. However, the extent to which spatially varying LAWT affects personalized mechanics-driven atrial flow predictions, and how these effects compare with plausible segmentation uncertainty, remains less quantified.

Here, we extended our multiscale myocardial mechanics modeling framework to incorporate spatially varying LAWT and voxel-level segmentation uncertainty, and quantified their impact on atrial wall mechanics and hemodynamics. Using gated CTA images, we constructed a baseline variable-thickness model of LA myocardium. We then generated geometric variants by perturbing the segmentation contours by 1-2 voxels to account for plausible inter-operator and image-driven uncertainty during model creation. For completeness, we also constructed a uniformly thick LA myocardium. We then applied a personalization workflow to the baseline case to estimate the multi-scale model parameters [22], and transferred these parameters across all thickness variants to isolate the effect of geometry from sensitivities in the material parameters and boundary conditions. Finally, we compared changes in regional deformation, stress measures, and clinically relevant flow metrics derived from moving-domain blood flow simulations. Collectively, these investigations advance our understanding of atrial biomechanics and hemodynamics, as well as the sensitivities of LAWT extraction.

## 2 Methods

The detailed multistage personalization workflow, including clinical data acquisition, model construction, boundary conditions, constitutive model, finite element methodology, and inverse analysis, is described in Shi et al. [22], and summarized in the Appendix (Fig. A1). A high-level overview of key components is provided below.

### 2.1 Myocardial Segmentation

#### Patient data

Ten equispaced, deidentified, gated CTA volumes spanning one cardiac cycle were obtained for a 50-year-old male subject with mild coronary artery disease from the database at the Veterans Affairs (VA) Healthcare System, Palo Alto, CA, USA, following IRB-approved protocol and HIPAA-compliant procedures. Based on the echo readings, the patient’s heart chambers were found to be normal in size, without wall motion abnormalities. Each CTA volume has a high spatial resolution (512 × 512 × 213 voxels) with a voxel size of 0.39 × 0.39 × 0.7mm^3^. More details about the imaging modality and CTA protocol are provided in Chen et al. [51].

#### LA myocardium with variable thickness (baseline)

We manually segmented the LA blood pool at the beginning of the reservoir phase in 3D Slicer (Fig. 1A, top-left). We then employed a modified version of the LAWT segmentation protocol to avoid ultrathin regions (i.e., sub-voxel resolution) and related numerical instabilities [22, 40]. Briefly, assuming comparable image intensities for the left atrial and ventricular myocardium, we used Seg3D2 [52] to expand the LA endocardial surface outward from the blood pool. We applied a four-point median filter to improve myocardial contrast. We then constructed a base layer of LA myocardium by subtracting the blood pool from a two-voxel dilation. We identified radiodensity bounds in the left ventricular myocardium and used them to guide subsequent morphological operations (Fig. 1A, top-right). Specifically, we dilated the base myocardial layer by 1 voxel, repeated this 3 times in succession, and refined it using a 2-voxel dilation-erosion sequence. Finally, we removed any residual mesh irregularities in MeshMixer (Autodesk Inc.). This myocardium served as the baseline reference model (Fig. 1A, bottom).

**Fig. 1:**
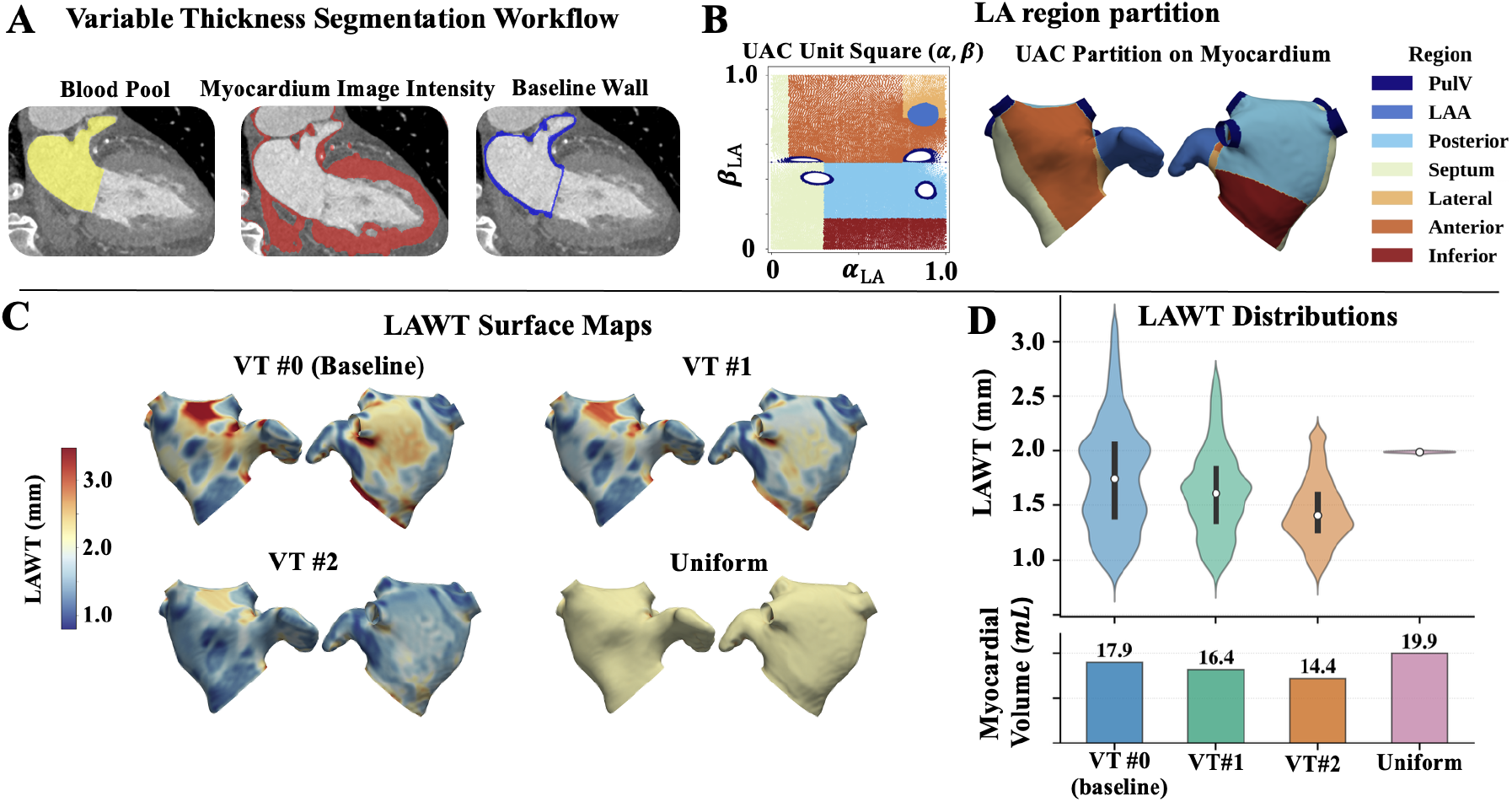
Extracting LA myocardium from CTA images with different wall-thickness sensitivities. (A) Representative slice illustrating the variable-thickness segmentation workflow, including (left) the segmented LA blood pool, (middle) the myocardium image-intensity range to guide LA wall segmentation, and (right) the final variable-thickness baseline myocardium after successive morphological operations and intensity-guided refinement. (B) LA is partitioned according to the Universal Atrial Coordinates (UAC) framework [54], with (left) regional partitioning on a 2D unit square that is independent of LA geometry, and (right) regional partitions mapped to the 3D surface. (C) LAWT surface maps for all the myocardial variants, including (top-left) baseline variable thickness model, VT#0, (top-right and bottom-left) two reduced morphological dilation variable thickness variants, VT#1 and VT#2, and (bottom-right) uniformly thick LA myocardium. (D) (top) Violin plots comparing LAWT distributions and (bottom) myocardial tissue volumes across the four models. The white circle denotes the median, the vertical bar spans the interquartile range (IQR), and the violin width represents the full distribution density. LA: left atrium; CTA: computed tomography angiography; PulV: pulmonary veins; LAA: left atrial appendage; LAWT: left atrial wall thickness; VT: variable thickness.

#### LA partitioning

To facilitate a local comparison of atrial wall mechanics and its relation with the local thickness, we partitioned the LA into seven non-overlapping anatomical regions: anterior, posterior, inferior, septum, lateral, pulmonary veins, and LAA (Fig. 1B) [46, 53]. To ensure a consistent, geometry-independent regional assignment across models, each surface was parameterized using the Universal Atrial Coordinates (UAC) framework introduced by Roney et al. [54]. Briefly, the pulmonary vein and LAA boundaries were first delineated to define anatomical landmarks and non-LA-wall regions. Three landmarks were then identified: one point each on the two superior pulmonary veins and one point at the fossa ovalis. Geodesic paths were used to define the lateral, septal, and LA roof boundaries. Laplace equations for two scalar fields (*α, β*) were then solved on the atrial surface with Dirichlet conditions prescribed on opposing anatomical boundaries. The resulting solutions were post-processed following the UAC framework to obtain the septal–lateral and anterior–posterior coordinates. *α* ranged between 0 and 1 from the septum to the lateral wall, while *β* ranged between 0 and 1 from the posterior mitral annulus through the roof to the anterior mitral annulus. The pulmonary veins and LAA were assigned directly from the predefined delineations. On the remaining LA wall, the septum and lateral regions were assigned first along the *α* extremes, with the septum defined as *α* < 0.30 for *β* < 0.5 and *α* < 0.10 for *β* ≥ 0.5, and the lateral region defined as *α* > 0.75 and *β* > 0.75. The residual surface was then divided by *β* into inferior (*β* < 0.175), posterior (0.175 ≤ *β* < 0.5), and anterior (*β* ≥ 0.5) regions (Fig. 1B).

#### LAWT computation

We computed local wall thickness using the Laplace-based field-line approach proposed in Bishop et al. [40]. To briefly summarize, we solve for the Laplacian of a scalar field,

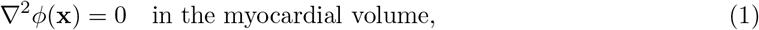

with Dirichlet conditions *ϕ* = 0 on the endocardium and *ϕ* = 1 on the epicardium, and no-flux conditions on the rest of the boundaries. A unit transmural vector is then defined as,

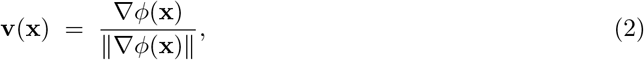

For each endocardial node **x**_0_, we trace a streamline **x**(*s*) starting at **x**_0_,

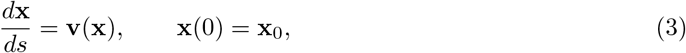

until the streamline reaches the epicardial surface at *s* = *L*(**x**_0_). The local wall thickness assigned to **x**_0_ is then the arc length of this streamline,

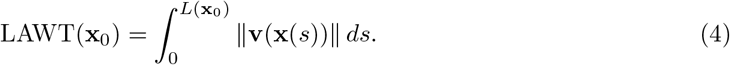

At a small minority of rim nodes (< 1.1% per model) where the gradient direction is tangent to a no-flux boundary and the streamline does not cross the wall, we instead assigned the Euclidean closest-point distance from the endocardial node to the epicardial surface as the measure of LAWT. A surface map of computed LAWT for the baseline model (VT#0) is shown in Fig. 1C (top-left).

#### LAWT variants

To represent plausible voxel-level segmentation uncertainty, we generated two additional LA myocardial variants by changing the base-layer dilation process. These two variants were created by dilating the base layer by 1 and 2 voxels, respectively, fewer than in the standard procedure, to account for plausible variations in image quality and inter-operator variability. Additionally, a uniformly thick LA myocardium was constructed by offsetting the endocardial surface outward along surface normals by 2mm [22]. Each perturbed surface was processed using the same smoothing, trimming, and meshing operations in MeshMixer (Autodesk Inc.) as those of the baseline model to remove self-intersections and ensure a smooth, watertight, thick-shelled volume. The spatial distributions of LAWT (Figs. 1C, D) highlight the degree of regional heterogeneity across all the variable-thickness cases. Myocardial volume comparisons suggest that the uniform case had the most tissue volume among the models considered here (Figs. 1D, bottom).

#### Fiber architecture

Left atrial fiber directions were computed using the rule-based approach described in detail in prior work [21, 22, 55]. Briefly, Laplace-Dirichlet solutions defined smoothly distributed regional and transmural coordinates. Fiber orientations were then computed as rotations within these coordinate systems to reproduce characteristic atrial fiber patterns that vary smoothly and continuously across the LA tissue [56–58]. All LAWT variants shared identical fiber-generation parameters, so any differences in the results could be attributed solely to thickness sensitivities.

### 2.2 Multiscale Mechanics

To ensure the physiological relevance of the boundary conditions, we adopted a multiscale model of left atrial mechanics in which the 3D finite element model of LA myocardium is coupled to a 0D lumped parameter network (LPN) model of the circulatory system via a robust, implicit, and modular coupling strategy [59]. Further, leveraging the subject’s multimodal clinical data, we adopted a three-step personalization strategy to efficiently estimate the multiscale model parameters that yield close agreement with clinical measurements when applied to the baseline LA model [18, 22]. This three-step personalization procedure is briefly described below, while the detailed workflow and personalization results are provided in the Appendix A.

#### 0D LPN parameter estimation

First, we replaced the 3D model with a 0D surrogate of the left atrium embedded within the closed-loop LPN-based circulation model. The LPN parameters were then optimized to match CTA-based cardiac chamber volumes and clinically measured and literature-informed pressures, resulting in a bespoke circulatory system replicating the patient’s physiology.

#### Passive mechanics personalization

Second, the LPN-predicted passive profile is used as input to iFEA for simultaneously estimating the stress-free configuration and myocardial passive mechanics parameters [21, 22]. The resulting Holzapfel–Ogden constitutive model parameters, together with the unloaded reference configuration, yielded reasonable agreement with image-based LA cavity volumes and endocardial displacements during passive expansion. This step was performed on the baseline variable thickness model (VT#0 in Fig. 1). The optimized baseline material parameters and the LPN-based pressure profile were consistently applied to all other LAWT variants (VT#1, VT#2, and Uniform in Fig. 1) to recover their respective stress-free geometries using the modified Sellier’s algorithm [21].

#### Contraction parameters estimation and multiscale FEA

Third, starting from the baseline stress-free configuration and iFEA-predicted passive material parameters, coupled with the personalized LPN model, the peak active stress magnitude and other active stress profile parameters are tuned until the simulated left atrial deformation and chamber volumes agree with the image data throughout the cardiac cycle [18, 22]. As in the previous step, the contraction parameters were estimated only for the baseline LAWT variant (VT#0) and consistently applied to all other cases.

### 2.3 Moving-domain Blood Flow Dynamics

The endocardial wall motion from the converged multiscale mechanics simulations was extracted and imposed as a moving-wall boundary condition to drive blood flow through the LA cavity. The fluid domain was constructed from the capped LA blood pool defined by the endocardial surface and the pulmonary vein and mitral valve annular openings. Further, the pressures at the pulmonary veins and flow rates through the mitral valve, predicted from the multiscale mechanics model, are applied as additional boundary conditions on these capped regions to drive atrial blood flow. Blood was modeled as an incompressible Newtonian fluid, and flow in the deforming LA cavity was solved using the arbitrary Lagrangian–Eulerian formulation of the Navier–Stokes equations. To maintain mesh quality during atrial deformation, the interior fluid mesh is smoothly morphed using displacements computed from a statics problem weighted by element Jacobians [60]. At the capped inlet and outlet planes whose motion was not explicitly available from the mechanics simulation, boundary velocities were projected from the surrounding tissue motion using an inverse-distance-weighting scheme [22]. Full details of the governing equations, boundary conditions, and numerical implementation are provided in Shi et al. [22].

### 2.4 Discretization, Numerical Methods, and Solver Parameters

The FEA-ready fluid and solid domains are discretized using four-noded tetrahedral elements generated using TetGen within SimVascular. We adopted the same meshing strategy and resolution based on a previously conducted mesh convergence analysis [22], resulting in approximately 150k elements in the tissue domain and 4.2M elements in the LA cavity for blood flow simulations. Variational multiscale stabilization is applied to both the systems of equations governing myocardial mechanics and blood flow to address incompressibility and convection-related instabilities, while allowing equal-order interpolation for velocity and pressure [22]. The resulting nonlinear system was solved using the Newton–Raphson method within a predictor–multi-corrector framework and integrated in time using the implicit generalized-*α* method. All FEA simulations were performed using our in-house multi-physics finite element solver, adapted from the open-source solver *svFSI* [61], and previously verified for cardiac mechanics and hemodynamics applications [18, 21, 22, 59, 62–67]. Further details on the choices of linear solver, preconditioners, and tolerances for passive mechanics, multiscale mechanics, and blood flow simulations are reported in detail in our prior work [22].

### 2.5 Mechanical and hemodynamic metrics of analysis

To quantify the mechanical and hemodynamic impact of LAWT approximation and segmentation differences, we computed various global and regional metrics from multiscale mechanics and blood flow simulations and correlated them against local myocardial thickness. These include LA cavity volumes, local measures of Cauchy stresses and strains, and endocardial pressure and shear environment, such as the instantaneous wall shear stress (WSS), time-averaged wall shear stress (TAWSS), and oscillatory shear index (OSI). All the metrics were computed over the last cardiac cycle after reaching a limit-cycle equilibrium. For each quantity assessed, we examined spatial distributions on the corresponding LAWT variant at key phases of the cardiac cycle. The summary statistics of various metrics include median, interquartile range (IQR), and upper percentiles. Region-wise analysis is performed using a 75th percentile pooled distribution threshold for the 1st principal of cycle-mean Cauchy stress, a 25th percentile pooled distribution threshold for TAWSS to characterize area under low WSS, and OSI greater than 0.3. We also quantified the effect size of each measure (thickness, Cauchy stress, TAWSS, and OSI) using Cliff’s *δ* [68], defined as,

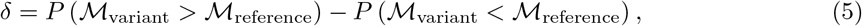

where positive values indicate larger metric values in the variant model and negative values indicate smaller values relative to VT#0 reference. Statistical significance was assessed using Mann-Whitney U tests with Holm correction for multiple comparisons.

## 3 Results

### 3.1 LAWT characteristics

The four myocardium models produced distinct LAWT distributions from the same CTA dataset (Fig. 1C-D). VT#0 had a broad, right-skewed distribution, with median thickness 1.78 ± 0.58mm, IQR 0.73mm, and myocardial volume 17.9 mL. The reduced-dilation variants were progressively thinner and more uniform: VT#1 had median thickness 1.62 ± 0.45mm, IQR 0.55mm, and volume 16.4 mL, whereas VT#2 had median thickness 1.41 ± 0.32mm, IQR 0.39mm, and volume 14.4 mL. The Uniform model had a narrowly distributed thickness of 1.99 ± 0.06mm, IQR 0.01mm, and the largest myocardial volume (19.9 mL). Overall, the myocardial volume ranged from 14.4 to 19.9 mL, a 38% difference due to variations in wall thickness.

### 3.2 Passive mechanics

Passive constitutive model parameters were optimized for VT#0 using iFEA [21] and mirrored to VT#1, VT#2, and the Uniform cases. The displacements required to recover the stress-free configuration from the initially segmented model were similar across the four LAWT variants (Fig. 2A–C). Median displacement ranged from 0.403 ± 0.148 cm in VT#0 to 0.416 ± 0.152 cm in VT#1, 0.434 ± 0.153 cm in VT#2, and 0.405 ± 0.151 cm in Uniform, respectively. Peak displacements approached 1 cm and were concentrated in the anterior region.

**Fig. 2:**
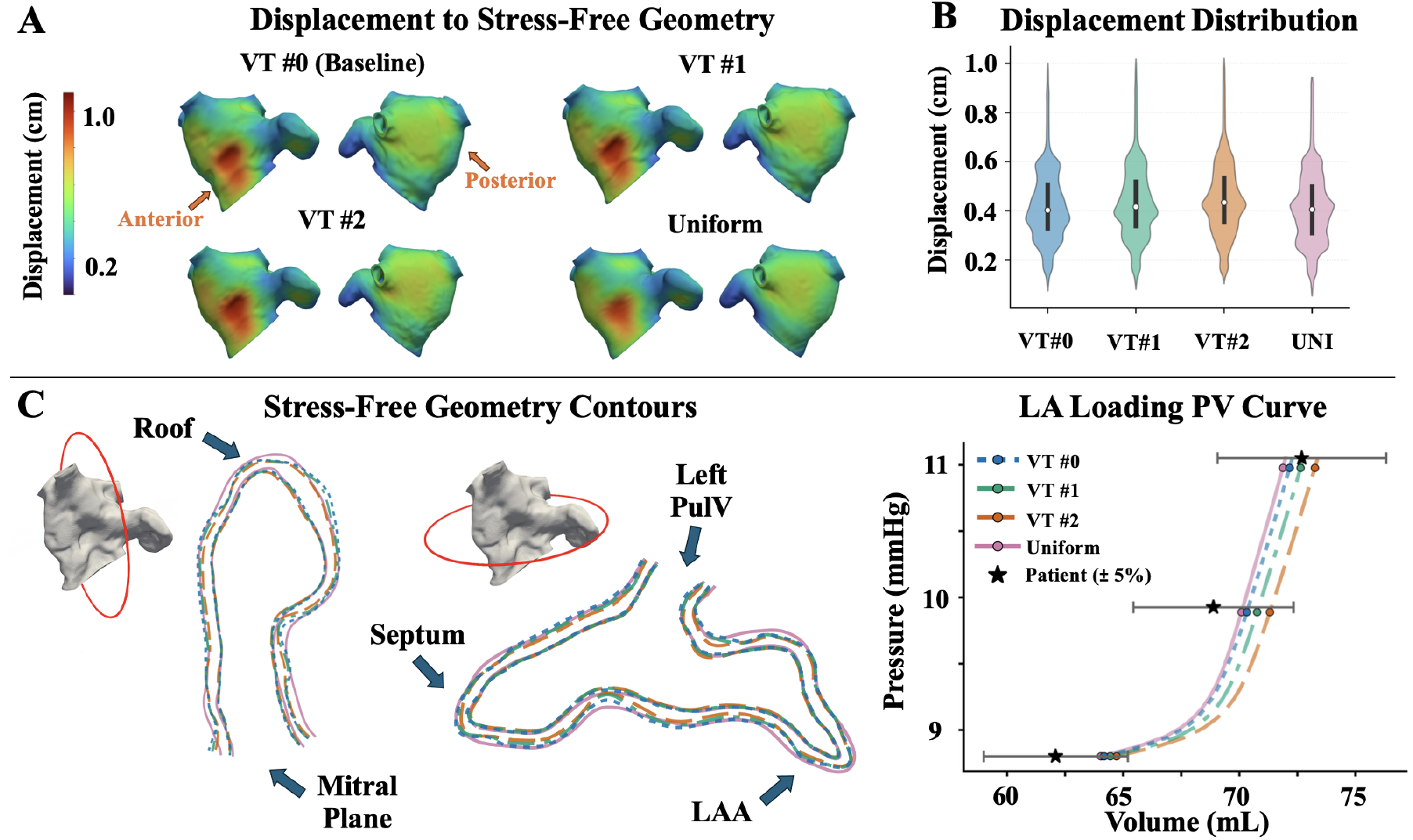
Effect of left atrial wall thickness (LAWT) on stress-free geometry recovery and passive pressure–volume response. (A) Surface maps of the displacement magnitude required to map each segmented myocardium from the imaged configuration to its stress-free geometry. For VT#1, VT#2, and Uniform, the stress-free configuration was recovered using Sellier’s method with passive material parameters identical to those optimized on VT#0 using iFEA [21]. (B) Violin plots summarizing displacement magnitude distributions; the white circle denotes the median and the vertical bar spans the interquartile range (IQR). (C) Surface contour comparisons from the recovered stress-free geometries for VT#0, VT#1, VT#2, and Uniform. The two contour overlays correspond to the cross-sectional slices indicated by the red ellipses on the 3D LA geometry. (D) LA pressure–volume relationship during passive loading for all four LAWT variants, with patient-derived target volumes shown with *±*5% uncertainty. LA: left atrium; LAA: left atrial appendage; PulV: pulmonary vein; iFEA: inverse finite element analysis; PV: pressure–volume; VT: variable thickness.

All four models reproduced the passive pressure–volume data within the prescribed ±5% uncertainty bounds at *P* = 8.8, 9.9, and 11.0mmHg (Fig. 2D). Compared to image data, VT#0 volume errors were +3.3%, +2.1%, and −0.7%. Among LAWT variants, the largest error was +4.2% in VT#2 at the lowest pressure; Uniform case errors were +3.1%, +1.7%, and −1.1%. Thinner variable-thickness walls produced larger cavity volumes at a given pressure, and the estimated unloaded volume ranged from 31.0 mL in VT#2 to 33.0 mL in Uniform.

### 3.3 Multiscale atrial mechanics

#### 3.3.1 LA volumes and pressures

All four models produced similar time-varying LA volume profiles, mostly within the prescribed ±5% uncertainty band compared to image-based volumes (Fig. 3a), and pressure–volume loops (Fig. 3b). The uniform-thickness case had the lowest minimum volume (35.6 mL) and largest LA stroke volume (39.1 mL). The variable-thickness models had higher minimum volumes (38.2–39.6 mL) and lower stroke volumes (36.8–37.4 mL), with the baseline VT#0 yielding a stroke volume of 37.3 mL. The corresponding LA ejection fractions were similar among the variable-thickness models (48.1–49.4%), whereas the Uniform model exhibited a higher LA ejection fraction of 52.3%. Total LA mechanical work, computed as the area enclosed by the pressure–volume loop, ranged from 7.1 to 8.1 mJ among variable-thickness models, with the baseline VT#0 expending 8.1 mJ, VT#1, 7.8 mJ, and VT#2, 7.1 mJ. The Uniform model performed the most total work (9.4 mJ, +16% to VT#0). This difference was driven almost entirely by the active booster-pump lobe (*A*-loop), which ranged from 3.5 mJ in VT#2 to 5.8 mJ in Uniform, whereas the reservoir lobe (*V* - loop) was nearly constant across all four variants (~3.6 mJ). Maximum chamber volume was similar across all four cases, and the largest differences occurred near the end of contraction and filling phases. Peak LA pressure ranged from 13.5mmHg in Uniform to 13.9mmHg in VT#0, and mean LA pressure ranged from 8.8 to 9.1mmHg.

**Fig. 3:**
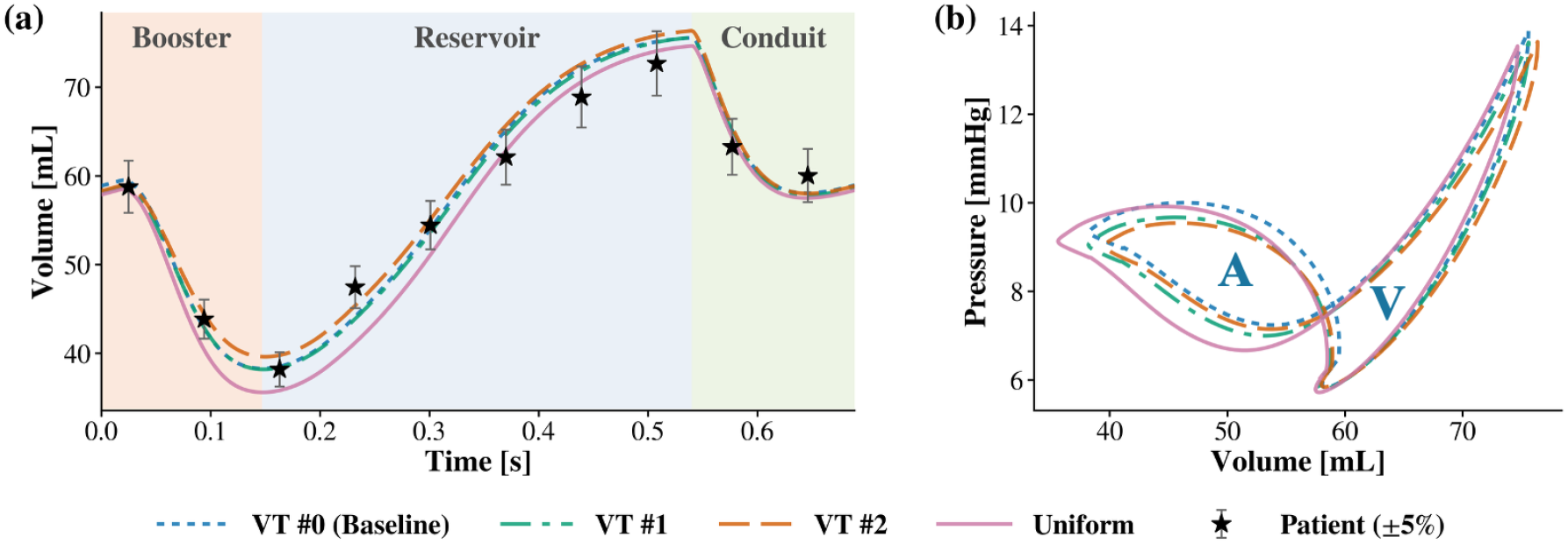
Effect of left atrial wall thickness (LAWT) variations on LA volume dynamics. (a) LA cavity volume over one cardiac cycle for the four LAWT variants, with image-based volumes shown with ±5% uncertainty. (b) Pressure–volume loop comparison over a cardiac cycle, highlighting the active (*A*-loop) and passive (*V* - loop) phases. All models used identical multiscale model parameters, including the passive and active material parameters and the same closed-loop loading framework, optimized on the VT#0 case.

#### 3.3.2 Circulatory system hemodynamics

Due to the closed-loop nature of the multiscale coupling between the 3D mechanics model and the 0D circulatory system model, we quantified differences in the response of the circulatory system across the LAWT variants (Appendix Fig. A2) and as percentage changes relative to the baseline VT#0 (Table 1). Cardiac output varied from 7.37 to 7.65 L/min. The left ventricular (LV) pressure–volume loops were closely clustered across all four cases (Appendix Fig. A2a). LV stroke volume ranged from 84.8 mL in VT#2 to 87.9 mL in VT#0, a spread of 3.6%, while LV ejection fraction ranged from 68.4 to 68.9% (Appendix Fig. A2a,d). The right ventricular (RV) PV loops showed relatively greater differences, with RV stroke volume approximately 9% higher in Uniform, VT#1, and VT#2 than in VT#0 (82.9 mL) (Appendix Fig. A2b,d). Peak LV pressure ranged from 105.5mmHg in VT#2 (−4.6%) to 110.6mmHg in VT#0 (Appendix Fig. A2c). Systemic arterial systolic pressure showed similar variations as that of LV pressure (104.3–109.3mmHg), whereas venous pressures were nearly unchanged (Appendix Fig. A2e,f). Overall, most circulatory metrics varied within 5% of the baseline VT#0 model, with RV and RA stroke volume being the exception and differing by over 9% and 6% compared to the baseline case (Table 1).

**Table 1:**
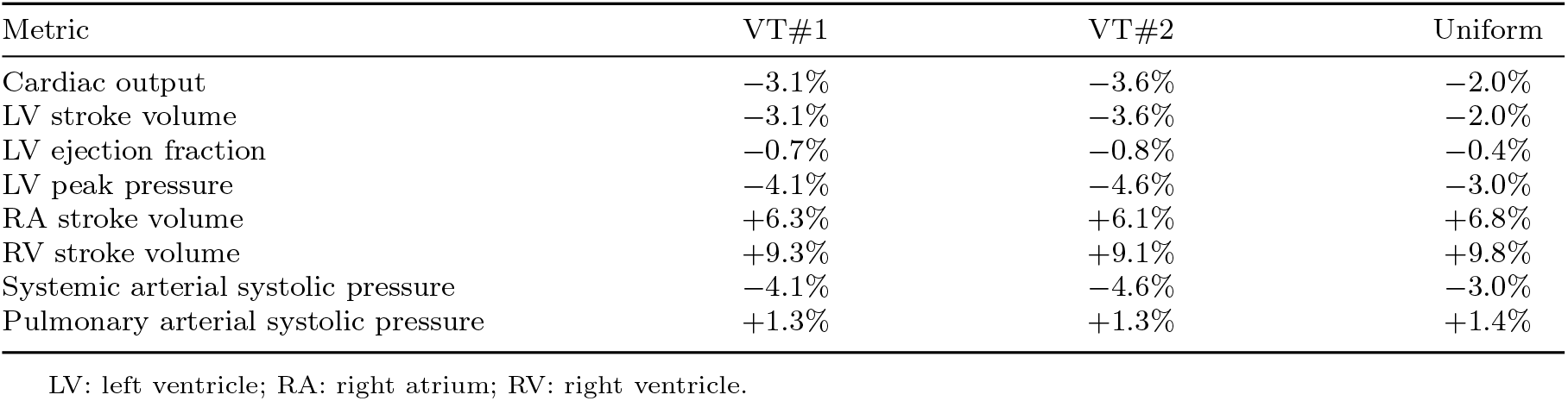
Percentage difference of key circulatory metrics relative to the VT#0 baseline model.

| Metric | VT#1 | VT#2 | Uniform |
| --- | --- | --- | --- |
| Cardiac output | -3.1% | -3.6% | -2.0% |
| LV stroke volume | -3.1% | -3.6% | -2.0% |
| LV ejection fraction | -0.7% | -0.8% | -0.4% |
| LV peak pressure | -4.1% | -4.6% | -3.0% |
| RA stroke volume | +6.3% | +6.1% | +6.8% |
| RV stroke volume | +9.3% | +9.1% | +9.8% |
| Systemic arterial systolic pressure | -4.1% | -4.6% | -3.0% |
| Pulmonary arterial systolic pressure | +1.3% | +1.3% | +1.4% |

LV: left ventricle; RA: right atrium; RV: right ventricle.

#### 3.3.3 Instantaneous wall mechanics

Instantaneous wall mechanics were compared at four representative phases during the cardiac cycle: the beginning of contraction or end of conduit phase (*t* = 0 s); end of contraction immediately before filling (*t* = 0.17 s); mid-filling (reservoir) phase (*t* = 0.3 s); and the end of filling (*t* = 0.52 s). Across these phases, internal stresses showed greater sensitivity to wall thickness variation compared to strains or displacements (Appendix Figs. A3–A8).

The left atrial myocardial displacements were less sensitive to thickness representation. At the end of filling (*t* = 0.52 s), mean displacements ranged from 6.10mm in VT#0 to 6.44mm (+5.47%) in VT#2; at all other phases, each variant’s mean displacement differed from that of the baseline VT#0 by no more than 6.3%. The largest displacements occurred along the anterior wall at the end of filling (*t* = 0.52 s, Appendix Fig. A3), followed by the posterior segment (Appendix Fig. A4). Endocardial displacements at all other phases remained similar qualitatively and quantitatively across all cases.

A comparison of the 1st principal strain showed only subtle differences between the four cases (Appendix Figs. A5-A6). Regions of larger strain were found along the lateral wall at the end of filling (*t* = 0.52 s), while strain patterns remained similar at all other phases of the cycle. At the end of filling (*t* = 0.52 s), the mean strain was 0.256 ± 0.106 in the baseline VT#0, 0.260 ± 0.106 in VT#1 (+2%), 0.269 ± 0.108 in VT#2 (+5%), and 0.239 ± 0.1 in Uniform (−6.4%). At the end of contraction (*t* = 0.17 s), the mean strain ranged from 0.2 to 0.213 across the four LAWT variants.

Local atrial stress distribution correlated well with the strain distribution, with higher stresses observed at the end of the filling phase compared to all other phases of the cycle (*t* = 0.52 s, Appendix Figs. A7-A8). Further, stress patterns exhibited marked differences due to wall thickness sensitivities. In particular, a comparison of 1st principal Cauchy stress showed that at the end of filling (*t* = 0.52 s), the mean stress was 10.2 ± 9.5 kPa in VT#0, 10.7 ± 9.4 kPa in VT#1 (+5%), 12.1 ± 9.8 kPa in VT#2 (+19%), and 8.6 ± 7.3 kPa in Uniform (−16%). The same ordering appeared at the end of contraction (*t* = 0.17 s), with mean stresses of 5.6 ± 4.0 kPa in the baseline VT#0, 5.7 ± 4.0 kPa in VT#1 (+2%), 6.2 ± 4.0 kPa in VT#2 (+10%), and 5.2 ± 3.5 kPa in Uniform (−7%).

### 3.4 Instantaneous atrial hemodynamics

Moving-domain blood flow simulations were performed by imposing the multiscale mechanics-predicted endocardial surface motion as a constraint. LA cavity pressures were found to be less sensitive to wall thickness variations (within 0.4 - 1.3mmHg). On the other hand, noticeable differences were found in the WSS distribution between variable-thickness cases and the uniform thickness case at the end of the conduit and filling phase (Appendix Figs. A9-A12).

Wall thickness variations had a modest influence on intracavitary pressure distributions (Appendix Figs. A9–A10). Inter-model differences were limited to 0.5–2.1mmHg depending on the cardiac phase. Pressures were the highest at the end of the filling phase across all LAWT variants (*t* = 0.52 s, 13.6–14.1mmHg), whereas LA pressures were the lowest at the end of the conduit phase (*t* = 0 s, 4.3–5.6mmHg).

Instantaneous WSS magnitude showed a stronger local structure than cavity pressure due to thickness-related differences in the left atrial wall motion (Appendix Figs. A11-A12). WSS was highest at the start and end of contraction (*t* = 0.00 s and *t* = 0.17 s), with localized peaks near the posterior wall and pulmonary vein ostia. In contrast, WSS remained low in the LAA at the end of filling (*t* = 0.52 s) across all models, with mean values of 0.13–0.18 Pa. Among the thickness variants, the Uniform model showed elevated mean instantaneous WSS at the end of the conduit phase at 2.48 Pa (+42% to VT#0) while the mean for the three thickness variants ranged from 1.64–1.75 Pa; conversely, at the end of passive filling, the Uniform model showed lower mean instantaneous WSS at 0.23 Pa (−39% to VT#0) while the three thickness variants ranged from 0.31–0.37 Pa.

### 3.5 Time-averaged wall mechanics and shear metrics

Time-averaged mechanics and flow metrics showed distinct spatial patterns across the LA endocardium (Fig. 4 and Appendix Fig. A13). The 1st principal of cycle-mean Cauchy stress was highest in the lateral wall (regional mean 10.3–14.4 kPa across all thickness cases, roughly twice that of the remaining regions, 4–8 kPa), with an additional localized stress concentration near the LAA junction. Time-averaged strain varied less than the cycle-mean Cauchy stress across regions and thickness cases. The regional mean 1st principal strain, computed over all non-pulmonary venous (PulV) regions and thickness variants, was 0.20 ± 0.03 with a coefficient of variation (CoV) 15%, which is about half the relative spread of the corresponding cycle-mean Cauchy stress (6.96 ± 2.64 kPa, CoV 38%). Compared with the variable-thickness cases, the Uniform model exhibited consistently lower time-averaged wall stress in every region (18–34 % below the variable-thickness mean), most pronounced in the septum (−34 %), the posterior wall, and least in the LAA (−18 %). Spatially, a low-stress region on the mid-anterior wall was present in all four models, but extended further toward the mitral annulus in the Uniform model. The posterior wall exhibited a more pronounced low-stress band near the mitral annulus in the Uniform model than in the baseline VT#0. Similar trends were observed for time-averaged strain, which was on average 10.6 ± 8.2% lower in the Uniform model than in VT#0 across the six non-PulV regions; the reduction was concentrated in the anterior, posterior, septal, and inferior walls (12–18% lower) and negligible in the lateral wall and LAA.

**Fig. 4:**
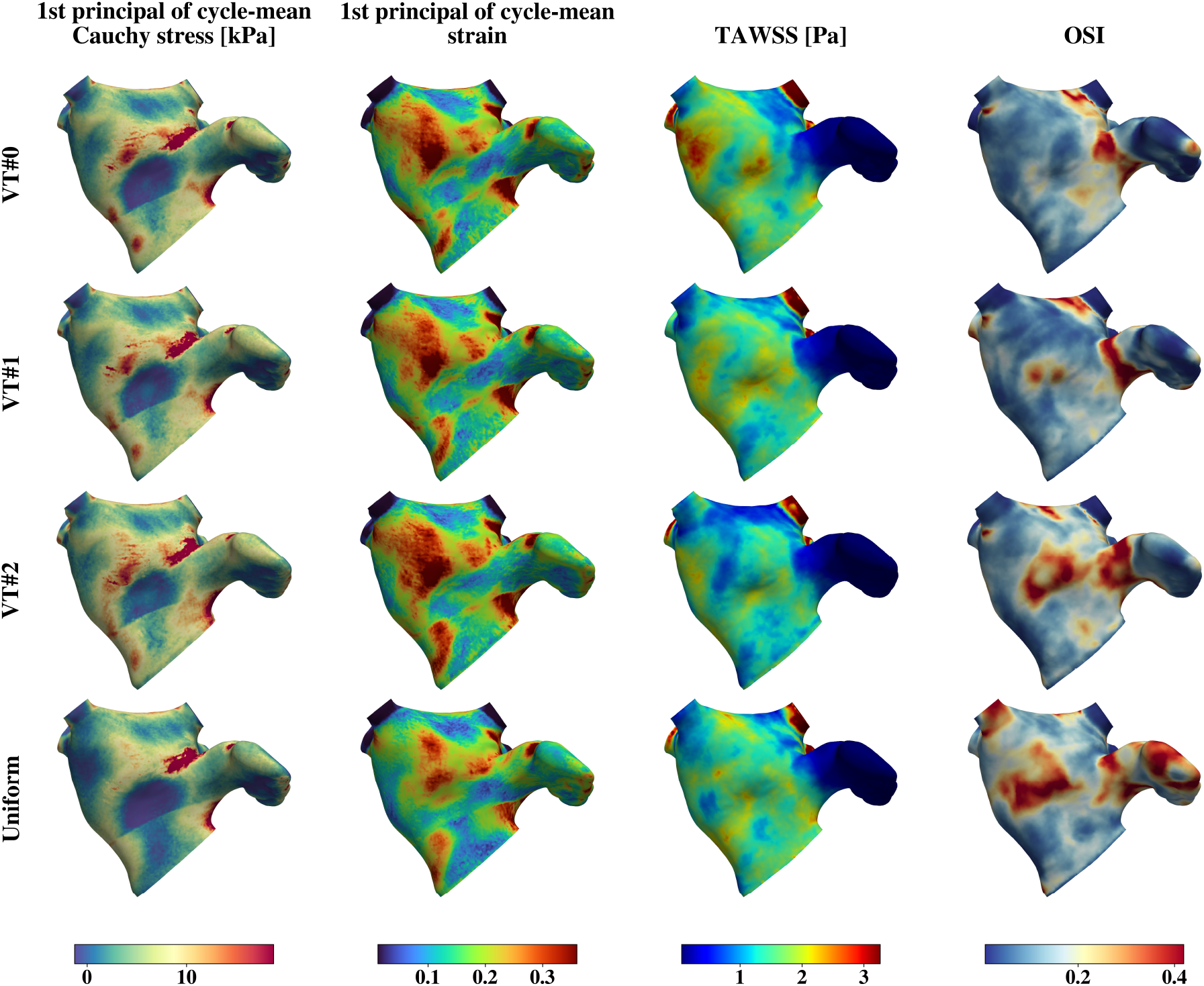
Comparison of time-averaged mechanics and hemodynamics metrics between LAWT variants along the anterior view. The same quantities are compared along the posterior view in Appendix Fig. A13. (column-wise) 1st principal Cauchy stress, 1st principal strain, TAWSS, and OSI. (row-wise) LAWT variants including VT#0 (baseline), VT#1, VT#2, and Uniform. LAWT: left atrial wall thickness; TAWSS: time-averaged wall shear stress; OSI: oscillatory shear index.

TAWSS was lowest in the LAA across all cases, while elevated OSI was concentrated in the LAA and the adjacent LA wall. Across the entire LA endocardium, mean TAWSS ranged from 1.16 Pa in VT#2 to 1.36 Pa in VT#0. The surface area fraction with TAWSS < 0.8 Pa (threshold based on the 25th percentile pooled distribution) remained similar across models (22.9–29.5%). In contrast, OSI showed larger inter-model differences. Mean endocardial OSI increased from 0.138 in VT#0 to 0.149 in VT#1, 0.164 in VT#2, and 0.188 in Uniform. The endocardial area fraction with OSI > 0.3 increased from 6.3% in the baseline VT#0 to 10.7% in VT#1, 14.1% in VT#2, and 19.0% in Uniform.

### 3.6 Wall thickness sensitivities on regional stress and shear measures

Region-wise effect sizes, computed using Eq. 5, confirmed that wall-thickness effects were spatially nonuniform (Fig. 5a). VT#1 and VT#2 had lower wall thickness than VT#0 across all regions, with larger reductions in VT#2. The uniform-thickness case had a higher wall thickness than VT#0 in the pulmonary vein, LAA, posterior, septal, anterior, and inferior regions, but a lower thickness in the lateral region.

**Fig. 5:**
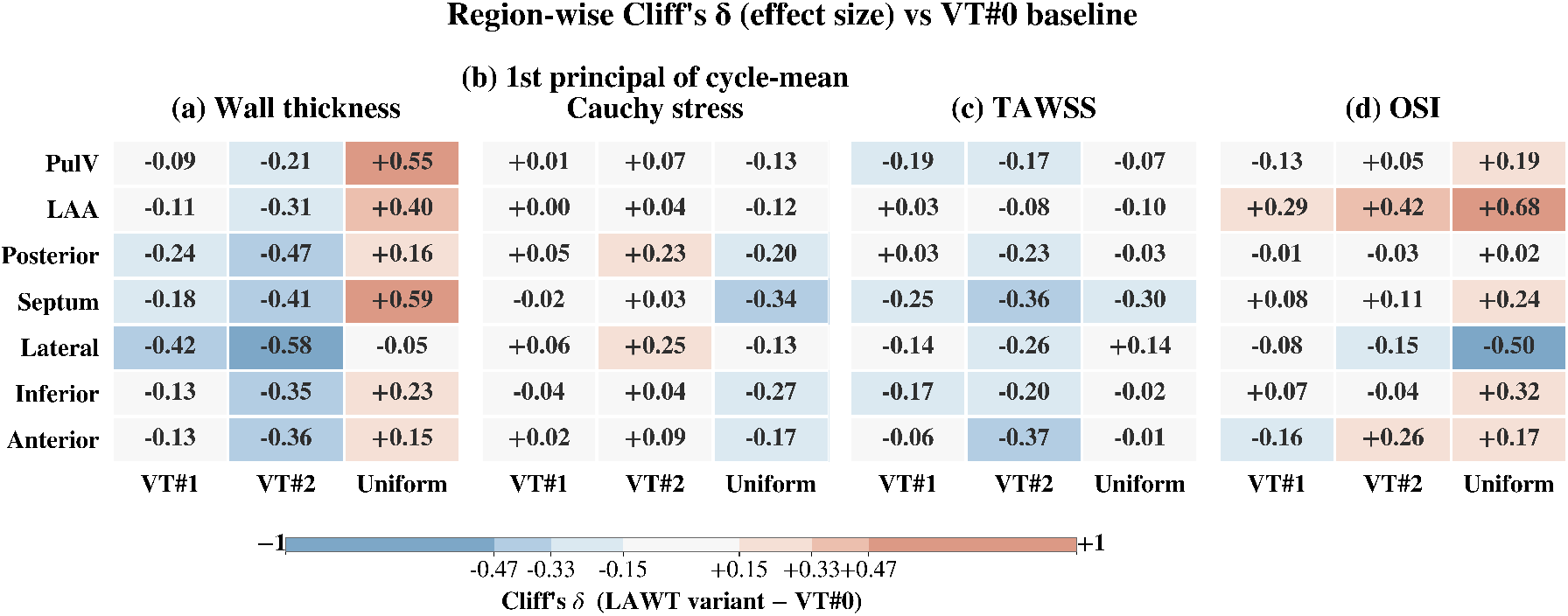
Region-wise Cliff’s *δ* values for each LAWT variant relative to the baseline VT#0 model. The metrics analyzed include (a) wall thickness, (b) 1st principal cycle-mean Cauchy stress, (c) TAWSS, and (d) OSI. Rows correspond to the LA regional partitions, including pulmonary veins (PulV), left atrial appendage (LAA), posterior, septal, lateral, inferior, and anterior regions. Positive values indicate larger values in the variant model than in VT#0, and negative values indicate smaller values. Hatched cells denote comparisons that were not significant after Holm correction (*p* > 0.05). LAWT: left atrial wall thickness; TAWSS: time-averaged wall shear stress; OSI: oscillatory shear index.

Region-wise effect sizes for the 1st principal Cauchy stress were small across all variants (Fig. 5b). Relative to the baseline VT#0, the uniform thickness model showed lower stress in all regions, whereas the thinner variable-thickness cases showed slightly higher stress, particularly in VT#2, while VT#1 remained largely unchanged.

Interestingly, both the variable-thickness cases and the uniform-thickness case exhibited a negative effect size for TAWSS across most regions (Fig. 5c). An exception is the lateral wall of the uniformly thick myocardium, which is subjected to higher TAWSS than in the baseline case. The same lateral wall, however, had a negative effect size for OSI relative to the baseline. All other regional segments of the Uniform case (i.e., PulV, LAA, posterior, anterior, inferior, and septal) showed a positive effect size for OSI with respect to the baseline VT#0, with the largest effect in the LAA (Fig. 5d).

Regional analysis revealed additional interesting trends in the LA mechanical response and shear dynamics (Fig. 6). Segmentally, the uniformly thick LA had the lowest fraction of high Cauchy stress. Among the variable-thickness variants, VT#2 has the highest fraction of high-stress regions compared to VT#0 and VT#1. Further, the lateral segment has the largest area coverage of higher stress among the entire LA, spanning 64–87% depending on the thickness distribution (Fig. 6, left).

**Fig. 6:**
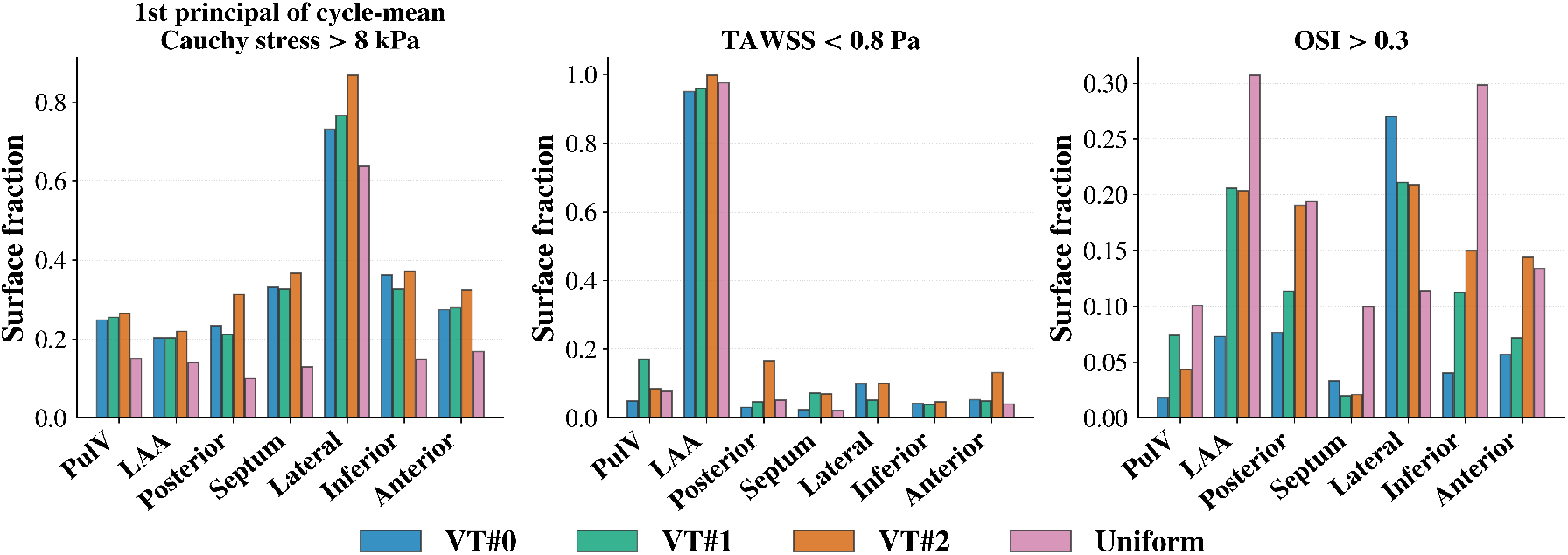
Comparison of region-wise surface-area fraction between the four LAWT variants constrained to meet a certain criterion, including (left) 1st principal cycle-mean Cauchy stress > 8 kPa, (center) TAWSS < 0.8 Pa, and (right) OSI > 0.3. PulV: pulmonary veins; LAA: left atrial appendage; LAWT: left atrial wall thickness; TAWSS: time-averaged wall shear stress; OSI: oscillatory shear index.

The LAA remained a low-shear region in every model: mean LAA TAWSS was 0.24–0.27 Pa, and 94.6–99.3% of the appendage surface had TAWSS < 0.8 Pa. The baseline VT#0 had the lowest low-shear fractions in the pulmonary veins, LAA, and posterior segments, while Uniform in the septum, lateral, inferior, and anterior segments. While VT#2 had a greater extent of low shear on the posterior, inferior, lateral, and anterior segments, VT#1 had a relatively higher fraction of low shear in the pulmonary veins (Fig. 6, center). Nevertheless, apart from LAA, all other LA segments had at most 20% fractional area under low shear stress (TAWSS < 0.8 Pa).

The uniformly thick LA showed the greatest span of high-OSI in most segments (PulV, LAA, septum, inferior, and posterior), except for the lateral wall (greatest in VT#0) and the anterior wall (greatest in VT#2). Among the variable-thickness cases, VT#2 generally had a greater extent of high-OSI than VT#0 and VT#1 (posterior, inferior, and anterior). The baseline VT#0 had the lowest span of high-OSI in most segments (PulV, LAA, anterior, posterior, and inferior) (Fig. 6, right). Lastly, mean LAA OSI increased from 0.115 in VT#0 to 0.173 in VT#1, 0.189 in VT#2, and 0.247 in Uniform, and the LAA surface fraction with OSI > 0.3 was 7.2% in VT#0, 20.4% in VT#1, 20.2% in VT#2, and 30.7% in Uniform.

## 4 Discussion

In this study, we evaluated how LAWT representation affects the left atrial biomechanical and hemodynamic function using a personalized multiscale modeling framework. Building on our recently developed, coupled 3D–0D mechanics-to-flow pipeline [18, 22], we compared four LA myocardium models constructed from the same gated CTA dataset: a baseline variable-thickness model, two reduced-dilation variable-thickness variants representing segmentation sensitivities, and the commonly employed uniform-thickness model. A key feature of our work is that the baseline multiscale mechanics model was personalized to patient data before assessing thickness sensitivities by mirroring model parameters and boundary conditions. We estimated the passive material parameters and the unloaded configuration on the baseline variable-thickness model using iFEA [21]. We then personalized the active mechanics parameters to reproduce full-cycle chamber-volume data. We then simulated atrial blood flow using the predicted wall motion while applying personalized boundary conditions, including pulmonary venous pressures and mitral flow rates. This design extends the previous uniform-thickness workflow [22] and enables us to systematically evaluate the effect of wall-thickness representation while minimizing confounding from differences in material parameters, loading conditions, and boundary conditions.

Assuming that the baseline LAWT extraction protocol yields a physiological representation of LA myocardium, we demonstrated that a uniformly thick myocardium overestimates thickness across most of the LA, except for the lateral wall (Fig. 5a, Sec. 3.6), leading to an overall increase in the tissue volume (Fig. 1D, Sec. 3.1). This increased myocardial mass will allow the uniformly thick LA to exhibit increased contractility (i.e., increased stroke volume and ejection fraction, Fig. 3, Sec. 3.3.1) and perform more mechanical work, while consistently maintaining lower internal stresses and strains, whether assessed instantaneously (Sec. 3.3.3) or when time-averaged over the cardiac cycle (Figs. 4, 5b, Sec. 3.5). We also showed that the thicker segments of LA had lower internal stress than the thinner segments, in accordance with Laplace’s law (Fig. 5a,b).

On the other hand, hemodynamic shear indices exhibited a more complex relationship with LAWT. Compared with myocardial mechanics, shear trends varied dynamically throughout the cardiac cycle relative to the baseline model (Sec. 3.4). The time-averaged shear, however, remained mostly lower for the uniformly thick myocardium than the baseline case (Fig. 5c), although the mean endocardial shear was less sensitive to thickness (Sec. 3.5). Conversely, OSI was substantially higher in regions with low shear for the uniform case compared to the baseline (Fig. 5d). The uniformly thick LA had the most area exposed to elevated OSI (~30%), while the baseline variable thickness model had the lowest exposure to high OSI (~7%). Fractional area analysis across LA segments showed that the LAA remained consistently low-shear in all cases, with over 90% of LAA exposed to low-TAWSS (Fig. 6)). The extent of low-TAWSS surface remained below 20% across all other LA segments and comparable across models.

The non-uniformly thick LA variants generally resulted in thinner myocardium than the baseline case (Fig. 5a), and therefore, were exposed to elevated internal stresses and strains (Fig. 5b, Secs. 3.3.3, 3.5, 3.6). Stroke volume, ejection fraction, and mechanical work were also generally lower (Fig. 3, Sec. 3.3.1). Mean shear was slightly lower while OSI was slightly higher over the entire surface (Sec. 3.5). Regionally, however, variably thick LA myocardium showed a mix of both high and low OSI relative to the baseline (Fig. 5d).

Previous LA modeling studies have shown that atrial wall thickness, anatomy, and material heterogeneity can affect different components of the mechanics and flow response. Augustin et al. [47] used electromechanical LA models to examine how local anatomy contributes to wall stress. In particular, it was demonstrated that the principal wall stress was associated with both wall thickness and curvature, with a stronger dependence on wall thickness during passive inflation. Feng et al. [34] developed a fluid–structure interaction model of LA with the mitral valve and evaluated how geometric modeling choices, including fiber architecture and wall-thickness representation, altered atrial mechanics and hemodynamics. Their results showed that uniform-thickness approximations changed atrial wall stress and strain, and that wall-thickness changes could also affect LAA flow velocity and pulmonary venous flow. More recently, Baptiste et al. [35] studied passive LA biomechanics in a cohort of patient-specific models and found that regional thickness is uncorrelated with local displacements but is strongly correlated with regional stiffness, when passive deformation was calibrated to image-derived data. Taken together, these studies suggest that certain mechanical and hemodynamic outputs are sensitive to LA wall thickness, especially local wall loading and flow features, while its effect on global function may be minimal when patient-specific material properties and physiological constraints are included. This provides the context for interpreting the present results, in which wall-thickness representation was evaluated leveraging personalized mechanics, closed-loop circulation, and moving-domain blood flow within a unified modeling framework.

Our results are broadly consistent with these prior thickness-sensitivity studies despite differences in methodology and problem setup. The largest differences were observed in the regional 1st principal Cauchy stress, especially at the end of filling. This trend is consistent with the mechanical expectation that, under comparable loading, changes in local wall thickness alter the amount and distribution of load-bearing tissue. It also agrees with Augustin et al. [47], who found an inverse correlation between wall stress and wall thickness during both passive inflation and active contraction, with a stronger association during passive inflation than during active contraction. Similarly, Feng et al. [34] reported lower maximum principal stress in uniform-thickness LA models than in the original patient-derived thickness model, with the lowest stress occurring in the thicker uniform-wall case, which aligns with our model predictions as well (Fig. 5, Sec. 3.6). Strains and displacements followed the same general ordering as stress in our models, but with lower relative differences (±5 −−6%). Thus, wall-thickness representation had a clearer effect on regional wall loading than on the kinematic measures used to match image-derived chamber motion. Shear analysis showed that OSI was most sensitive to thickness representation, particularly the uniform thickness model, which had over 30% of LAA exposed to high OSI. These results extend the flow sensitivity reported by Feng et al. [34] by separating shear magnitude from shear directionality, as a low-TAWSS criterion alone would make it less sensitive to thickness by yielding similar appendage shear exposure despite being consistently over 90%. With multiphysics approaches to modeling cardiac function being increasingly adopted, an unphysiological representation of LAWT may lead to inaccurate interpretation and classification of thrombogenic surrogates (areas with low shear and high OSI), particularly in the context of atrial fibrillation.

Several limitations and design choices must be considered when interpreting our results. The passive and active material parameters were personalized on the baseline VT#0 and applied ‘as-is’ to the remaining geometries. This was intentional because the objective was to isolate the downstream impact of wall-thickness representation under consistent material description and loading conditions. If the personalization steps, including iFEA and active mechanics characterization, were performed separately on each geometry, the optimizer would likely compensate for geometry-specific differences through material parameters. The segmentation variations were also limited to reduced-dilation variants of the baseline myocardium, which capture one plausible axis of voxel-level uncertainty but do not include localized segmentation errors near the LAA, pulmonary veins, or mitral plane. The underlying workflow also incorporated modeling choices described in detail in Shi et al. [22], including uniform material parameters, phenomenological active-stress model without excitation–contraction coupling, rule-based fiber architecture, simplified epicardial constraints, and spatially uniform mechanics pressure loading. Further, although we performed a controlled comparison of wall-thickness representations under matched physiological conditions, the baseline parameterization was based on a single patient’s data. However, the quantified sensitivities may vary across subjects with different chamber sizes, appendage morphologies, fibrosis burdens, rhythm states, or loading conditions. Finally, blood-flow simulations were intentionally driven by imposed endocardial wall motion from the mechanics model, which isolates the effect of mechanics-derived wall motion on flow; a fully coupled fluid–structure interaction formulation would account for feedback from intracavitary flow and local pressure variations, which is a subject for future investigation.

In summary, we leveraged a left atrial digital twinning framework to evaluate the sensitivity of myocardial mechanics and hemodynamics to local wall-thickness representation. The digital twin framework is based on our recently developed multiscale model of left atrial mechanics and blood flow personalized to a subject’s multimodal clinical data. We demonstrated that a uniform thickness assumption overestimates local thickness, leading to increased myocardial mass, stroke volume, and mechanical work, although overall displacements and strains were less sensitive to thickness. However, mean tissue stress had a larger coefficient of variation than strains. Further, oscillatory shear was found to be the most sensitive to wall thickness, particularly in the left atrial appendage, which may impact thrombogenic assessment under arrhythmic conditions.

## Abbreviations

AF: Atrial Fibrillation
CTA: Computed Tomography Angiography
iFEA: Inverse Finite Element Analysis
LA: Left Atrium
LAA: Left Atrial Appendage
LAWT: Left Atrial Wall Thickness
LPN: Lumped Parameter Network
OSI: Oscillatory Shear Index
TAWSS: Time-Averaged Wall Shear Stress
WSS: Wall Shear Stress

## Acknowledgments

BG and VV would like to acknowledge financial support from the American Heart Association’s Second Century Early Faculty Independence Award (#24SCEFIA1260268) and the US National Science Foundation’s CAREER Award (#2443726) in performing this work. BG and VV would also like to acknowledge partial funding support from Columbia University SEAS Interdisciplinary Research Seed (SIRS) grant / Blavatnik Fund. LS would like to acknowledge financial support from the US National Heart, Lung, and Blood Institute (#R15HL181637). Computing resources from ACCESS (SDSC Expanse) and the Columbia Ginsburg HPC cluster were used for this work.

## Declarations

### Human Subjects

This single-center study was performed following HIPAA-compliant processes and guidelines and was approved by the Columbia University Irving Medical Center (CUIMC) Institutional Review Board (IRB#AAAV3363) with a data use agreement with the Veterans Affairs (VA) Healthcare System, Palo Alto, CA, USA.

### Consent for Publication

All authors have agreed with the content and given explicit consent to submit the manuscript for publication.

### Data Availability

Data will be shared upon a reasonable request from the corresponding authors.

## Appendix A

**Appendix**

**Fig. A1:**
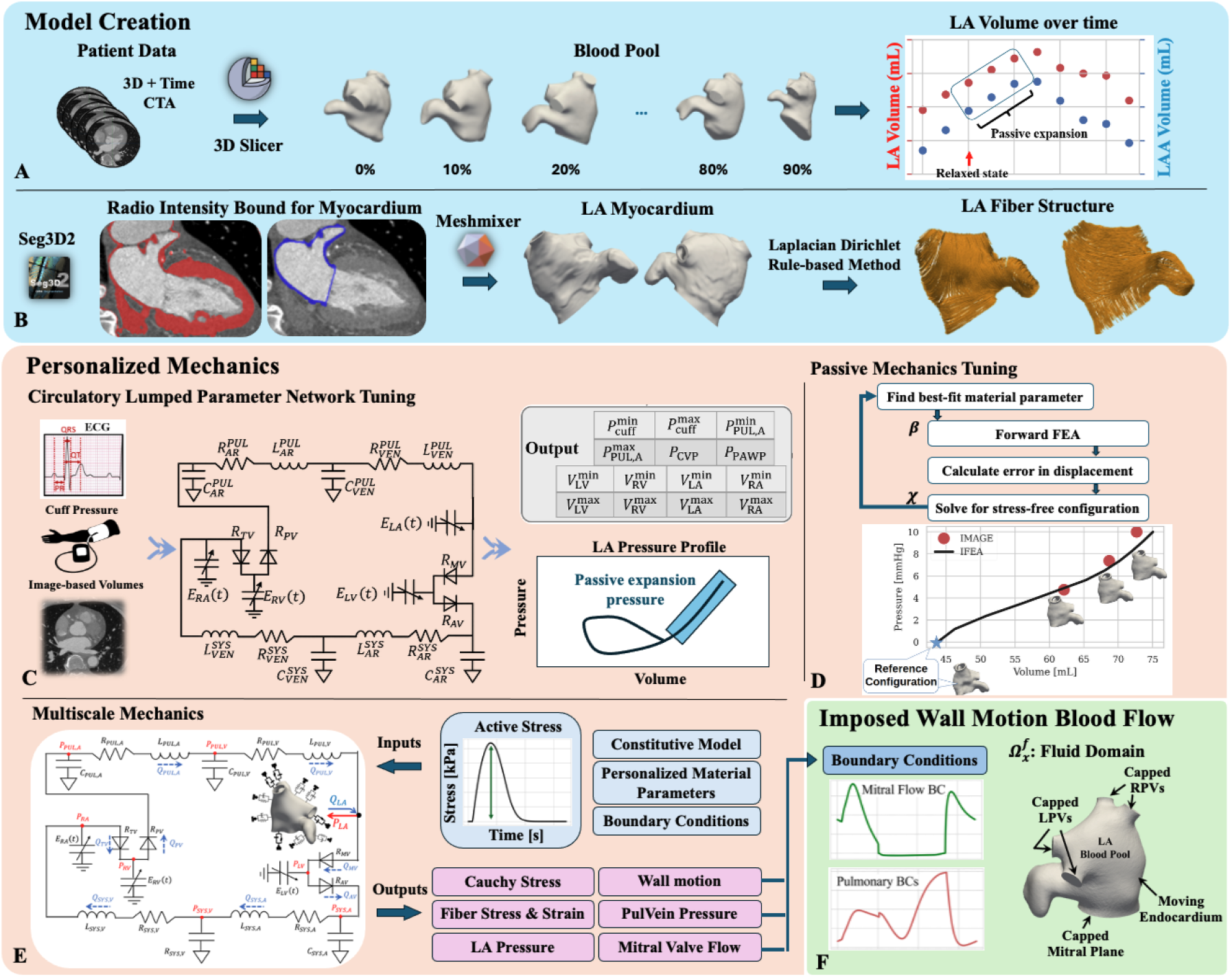
Overview of the personalized multiscale left atrial (LA) mechanics-to-flow pipeline used in this study. The top blue block summarizes model creation. (A) ECG-gated cardiac CTA provides time-resolved LA blood-pool geometries over the cardiac cycle, from which image-based chamber volumes are extracted to identify the passive expansion phase. (B) The LA myocardium is reconstructed from the segmented blood pool using intensity-guided morphology operations informed by left ventricular (LV) myocardium radiodensity bounds, followed by rule-based bilayer fiber assignment using a Laplacian–Dirichlet framework. In the present study, this model-creation procedure is used to generate the variable-thickness reference model, the uniform-thickness model, and the segmentation-uncertainty variants. The middle peach block summarizes personalized mechanics. (C) A closed-loop 0D lumped-parameter network (LPN) is tuned using ECG data, cuff pressure, and image-derived chamber volumes to obtain subject-specific circulatory parameters and the LA pressure waveform. (D) During passive filling, inverse finite element analysis is performed to identify the unloaded reference configuration and best-fit passive constitutive parameters by matching model-predicted deformation to image-derived motion. (E) The personalized passive model is coupled to the 0D circulation for whole-cycle multiscale mechanics simulations, in which active contraction parameters are tuned to reproduce subject-specific LA deformation and chamber volume dynamics; outputs include chamber pressure, wall stress/strain, wall motion, and valve/venous flow quantities. The right green block summarizes imposed-wall-motion blood-flow simulation. (F) The converged endocardial wall motion is prescribed as a moving boundary condition in an ALE simulation of the LA cavity, with pulmonary venous and mitral boundary conditions obtained from the coupled mechanics model, to quantify intra-atrial hemodynamics.

## A.1 Constitutive Model

We model the myocardium as a hyperelastic, nearly incompressible material, using the orthotropic modified Holzapfel–Ogden (HO) model. The isochoric components of the corresponding Gibbs free energies are given as,

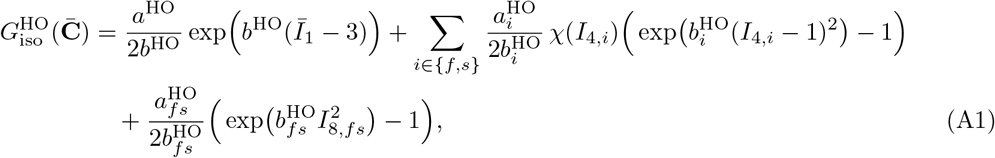

We also incorporate a smooth approximation of the Heaviside function,

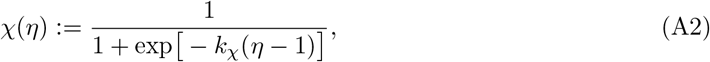

to avoid any numerical instabilities under small compressive strains.

In Eq. (A1), the set 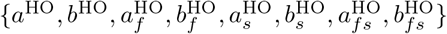 defines the parameters for the HO model. Further, 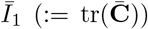 is the isotropic invariant that captures isochoric deformations, 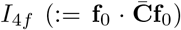 and 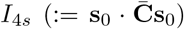 are the transverse invariants for the fiber and sheet directions, respectively, and 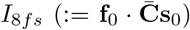 is the anisotropic invariant that captures the fiber–sheet interactions.

The pulmonary veins are modeled as neo-hookean material with elastic modulus of 3 × 10^6^ *dyne/cm*^2^ and Poisson ratio of 0.4833.

**Fig. A2:**
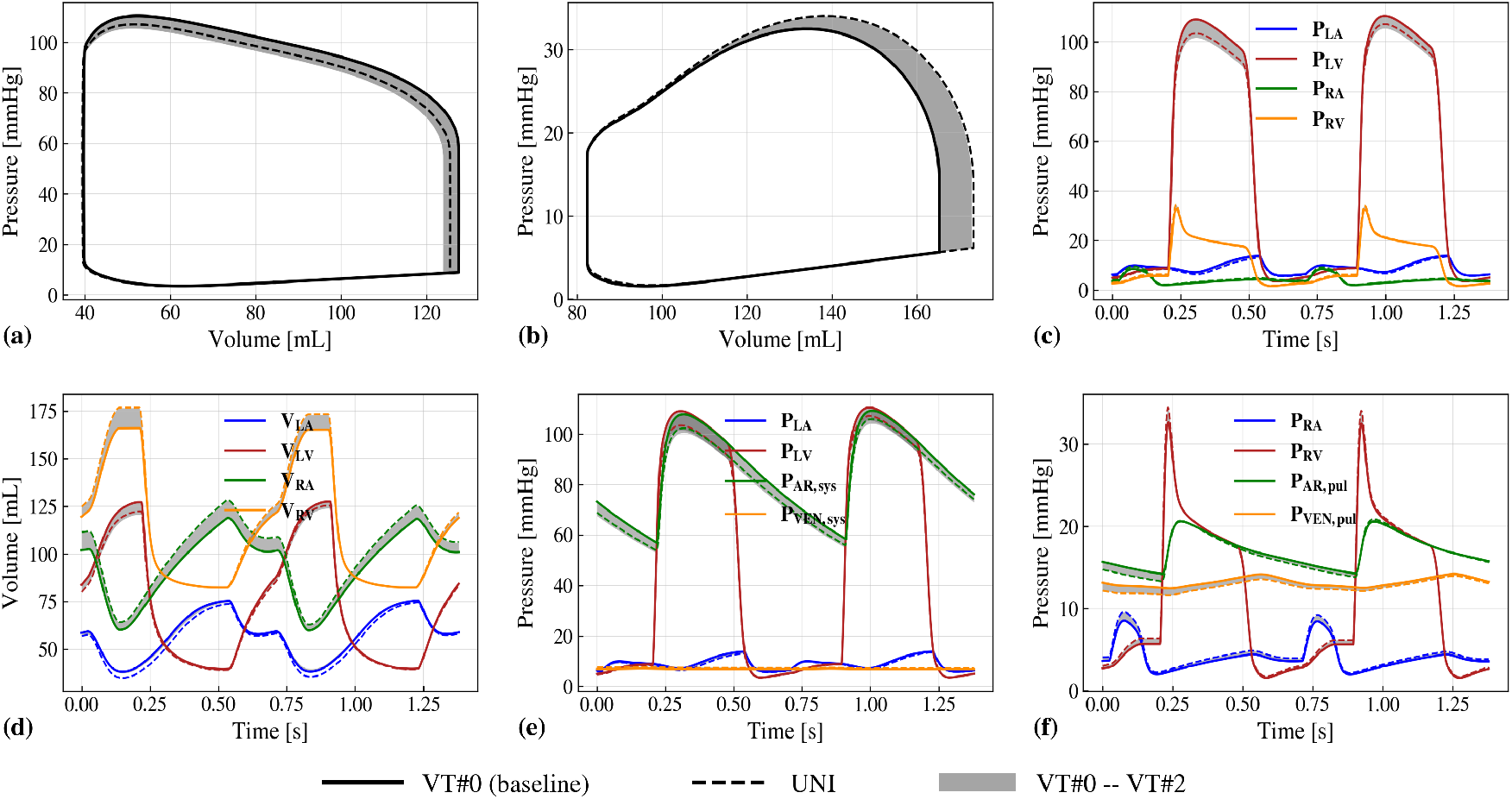
Effect of LAWT sensitivities on closed-loop circulatory hemodynamics from coupled 3D–0D multiscale mechanics simulations. (a, b) Pressure–volume loops of the left and right ventricles. (c, d) Cardiac chamber pressures and volumes over two cardiac cycles. (e) Left-side pressures, including LA, LV, systemic arterial, and systemic venous pressures. (f) Right-sided pressures, including RA, RV, pulmonary arterial, and pulmonary venous pressures. Solid lines denote VT#0; dashed lines denote Uniform; the shaded band denotes the range spanned by VT#1 and VT#2. LAWT: left atrial wall thickness; LA: left atrium; LV: left ventricle; RA: right atrium; RV: right ventricle; SYS/PUL AR/VEN: systemic/pulmonary arteries/veins.

**Fig. A3:**
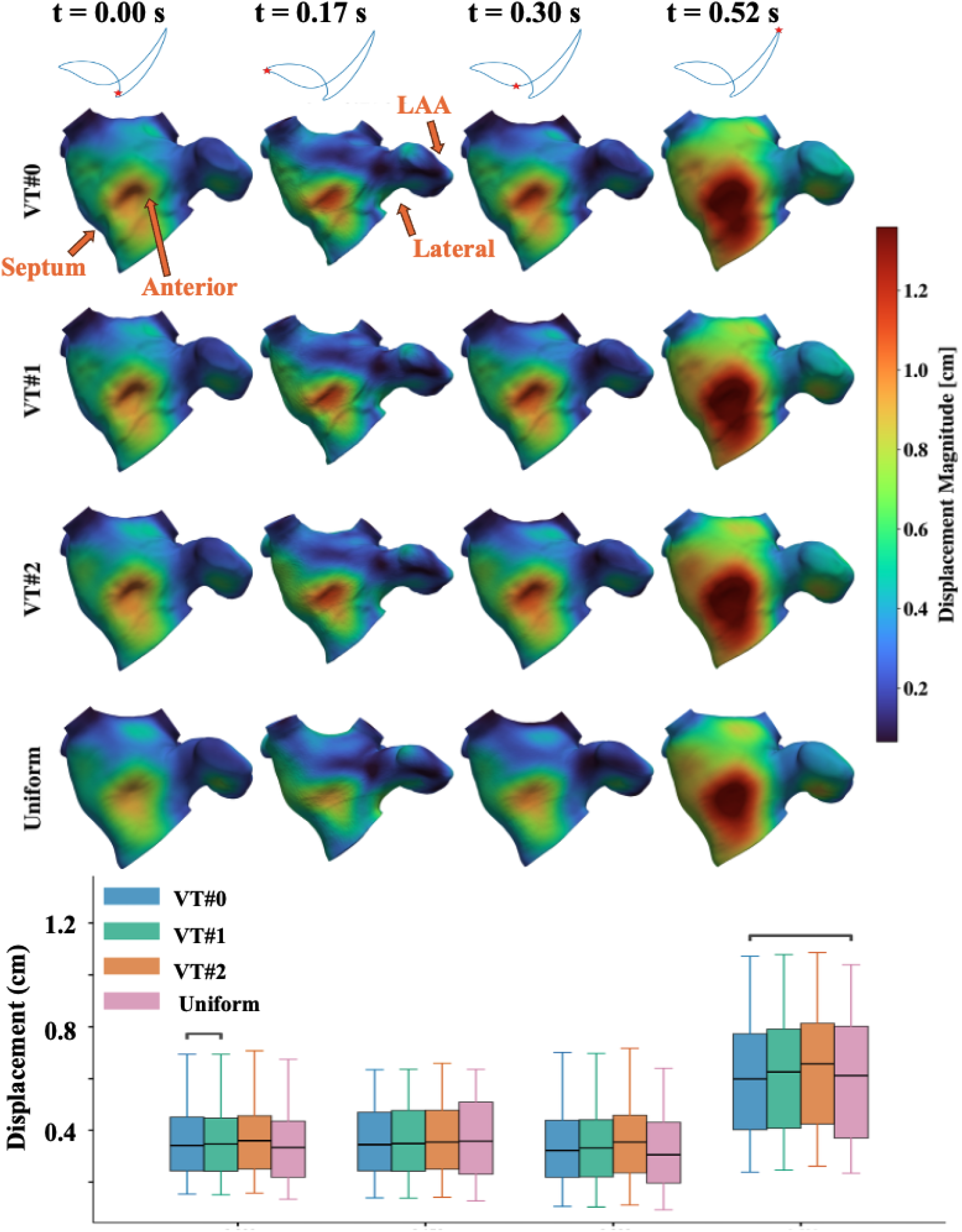
Comparison of instantaneous displacements between the four LAWT variants (row-wise) across key phases of the cardiac cycle (column-wise). (top) Surface map of the endocardial displacement field along the anterior view. (bottom) Grouped boxplots of nodal endocardial displacements at each phase, with whiskers spanning the 5th–95th percentile.

**Fig. A4:**
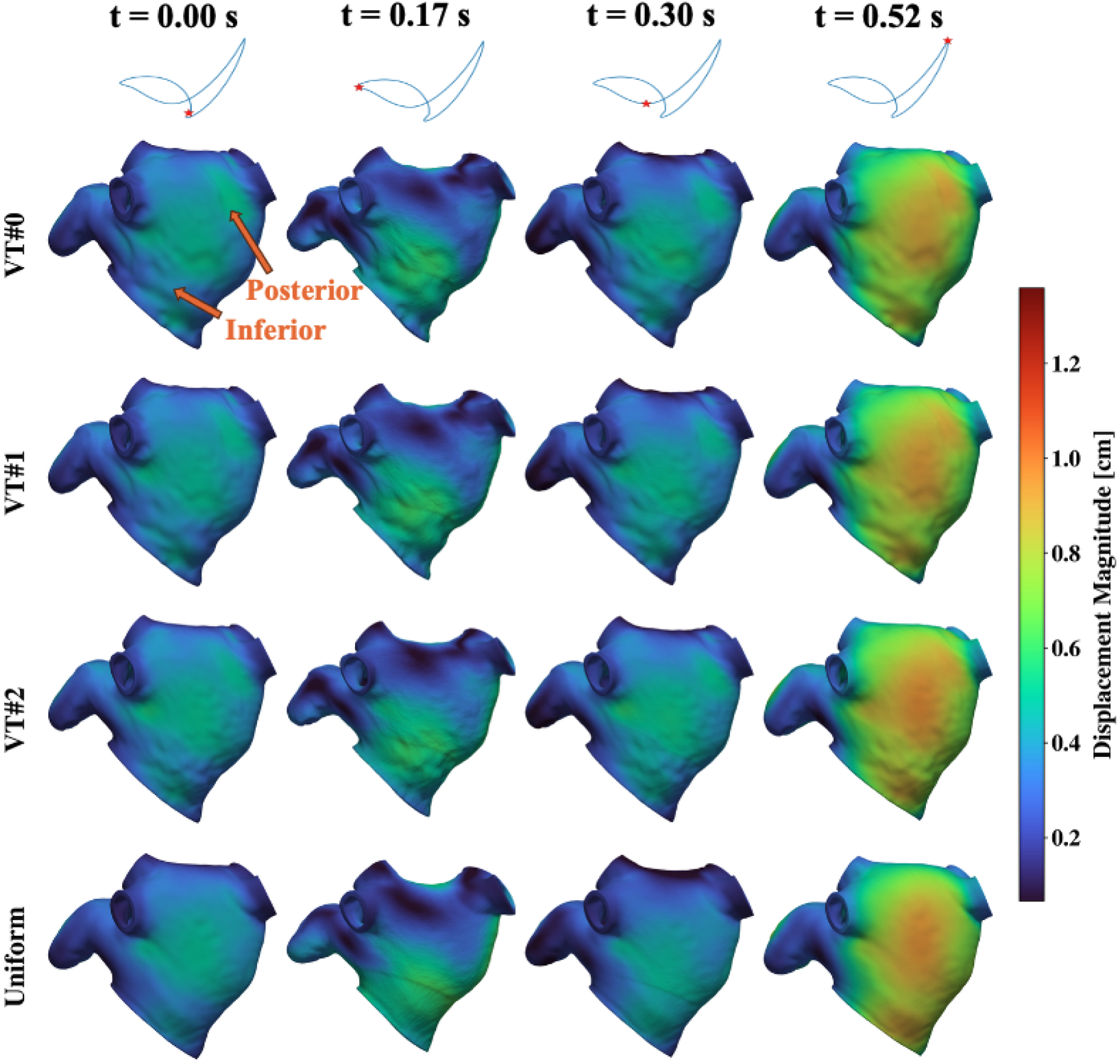
Comparison of instantaneous displacements between the four left atrial wall thickness variants (row-wise) across key phases of the cardiac cycle (column-wise) rendered along the posterior view.

**Fig. A5:**
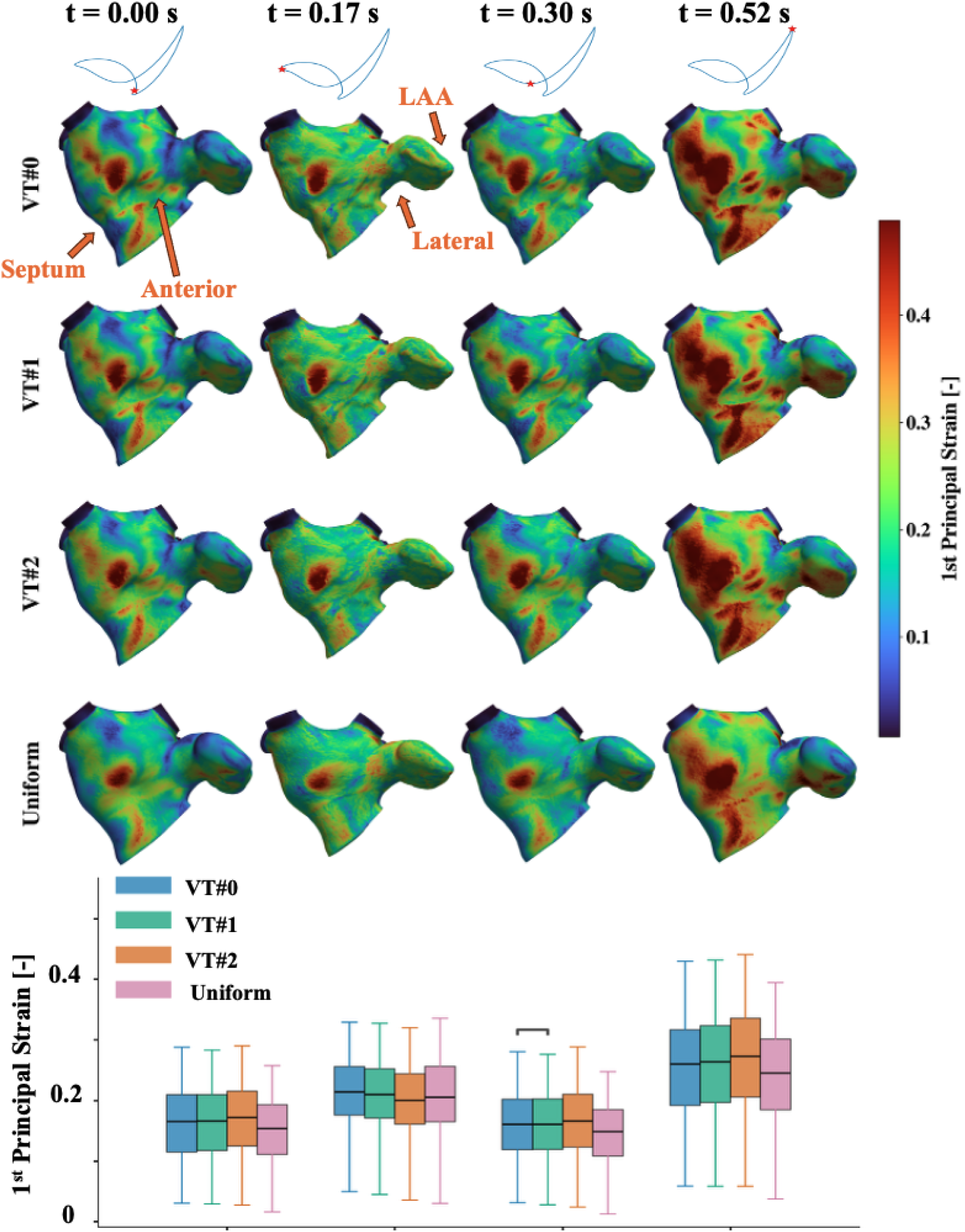
Comparison of instantaneous 1st principal strains between the four left atrial wall thickness variants (row-wise) across key phases of the cardiac cycle (column-wise). (top) Surface map of the endocardial strain field along the anterior view. (bottom) Grouped boxplots of nodal endocardial strains at each phase, with whiskers spanning the 5th–95th percentile.

**Fig. A6:**
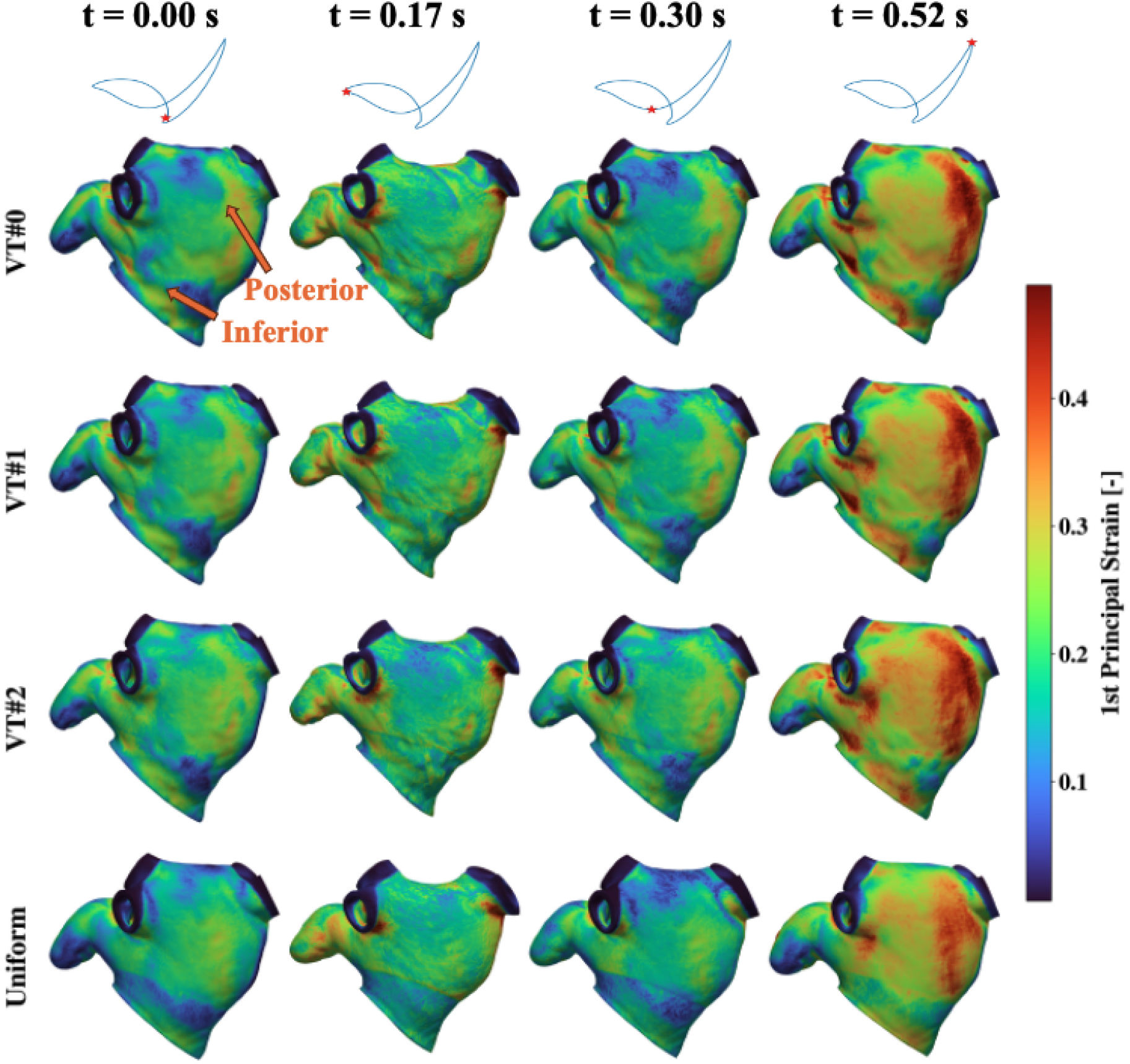
Comparison of instantaneous 1st principal strains between the four left atrial wall thickness variants (row-wise) across key phases of the cardiac cycle (column-wise) rendered along the posterior view.

**Fig. A7:**
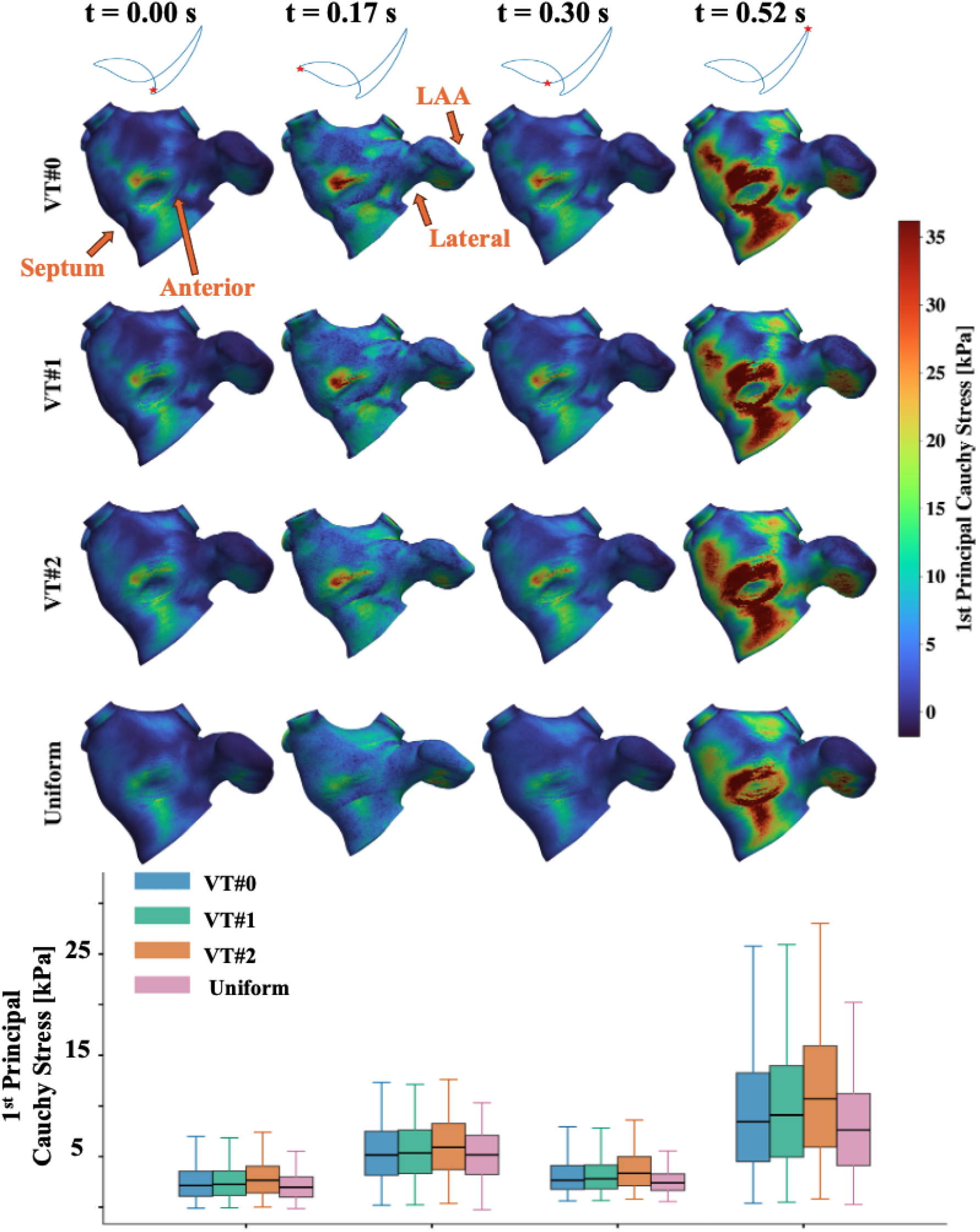
Comparison of instantaneous 1st principal Cauchy stress between the four left atrial wall thickness variants (row-wise) across key phases of the cardiac cycle (column-wise). (top) Surface map of the endocardial stress field along the anterior view. (bottom) Grouped boxplots of nodal endocardial stresses at each phase, with whiskers spanning the 5th–95th percentile.

**Fig. A8:**
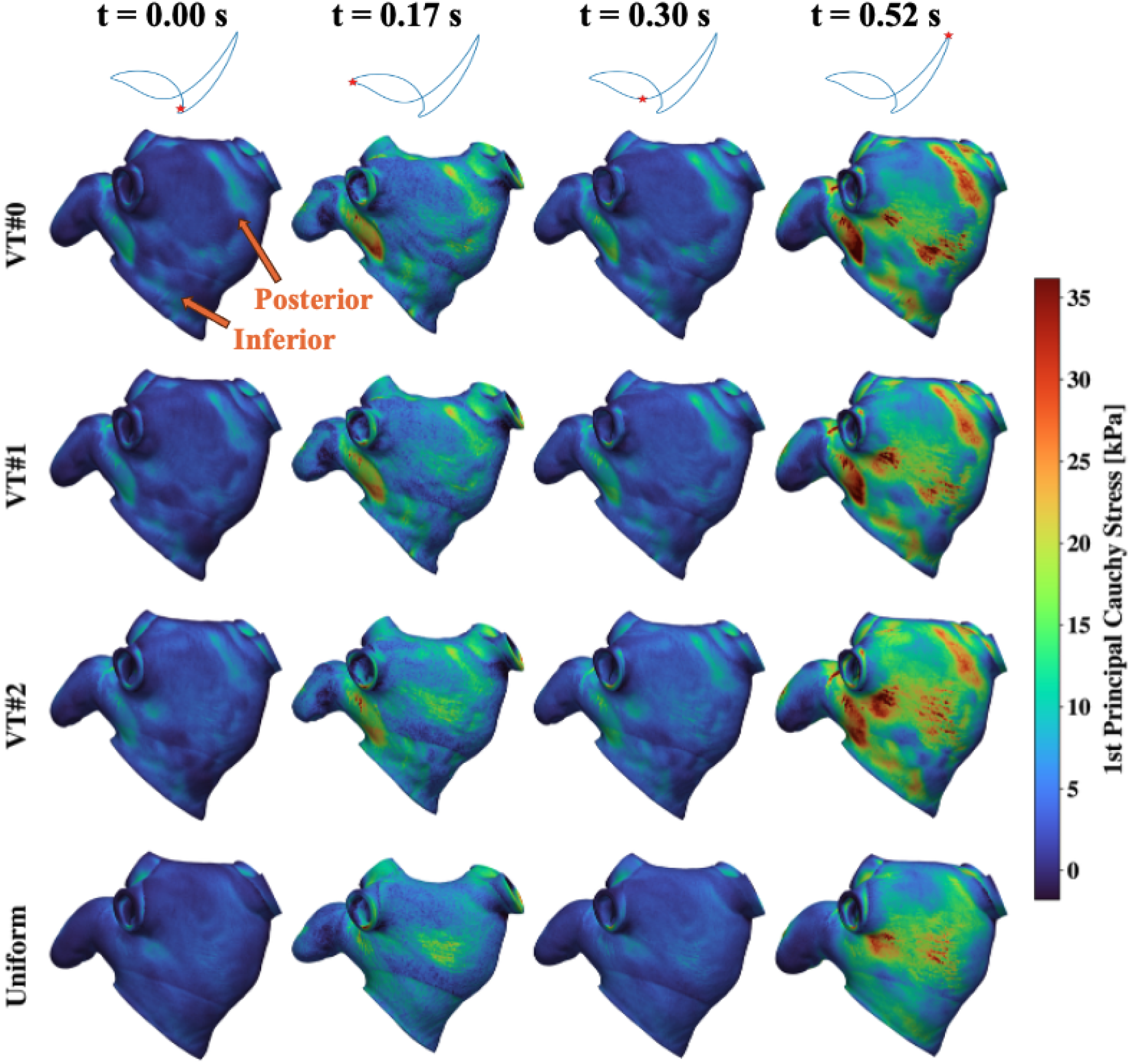
Comparison of instantaneous 1st principal Cauchy stress between the four left atrial wall thickness variants (row-wise) across key phases of the cardiac cycle (column-wise) rendered along the posterior view.

**Fig. A9:**
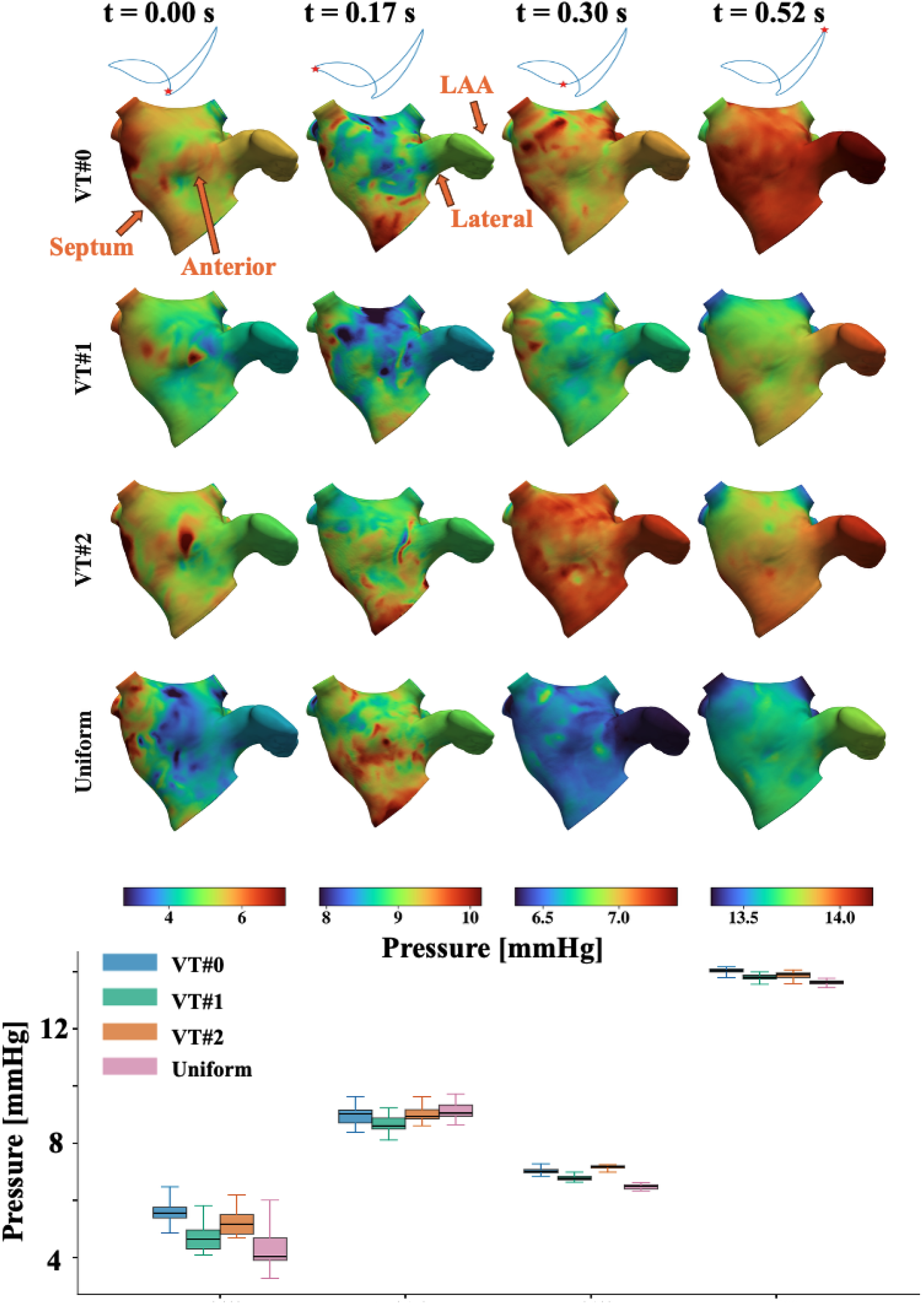
Instantaneous intracavitary pressure across the four LA models at four cardiac-cycle phases. Top: posterior surface maps with phase-specific colorbar ranges. Bottom: grouped boxplots of pressure of the LA endocardium surface at each phase, with whiskers spanning the 5th–95th percentile. Each phase uses a different pressure scale to show spatial variation within that phase.

**Fig. A10:**
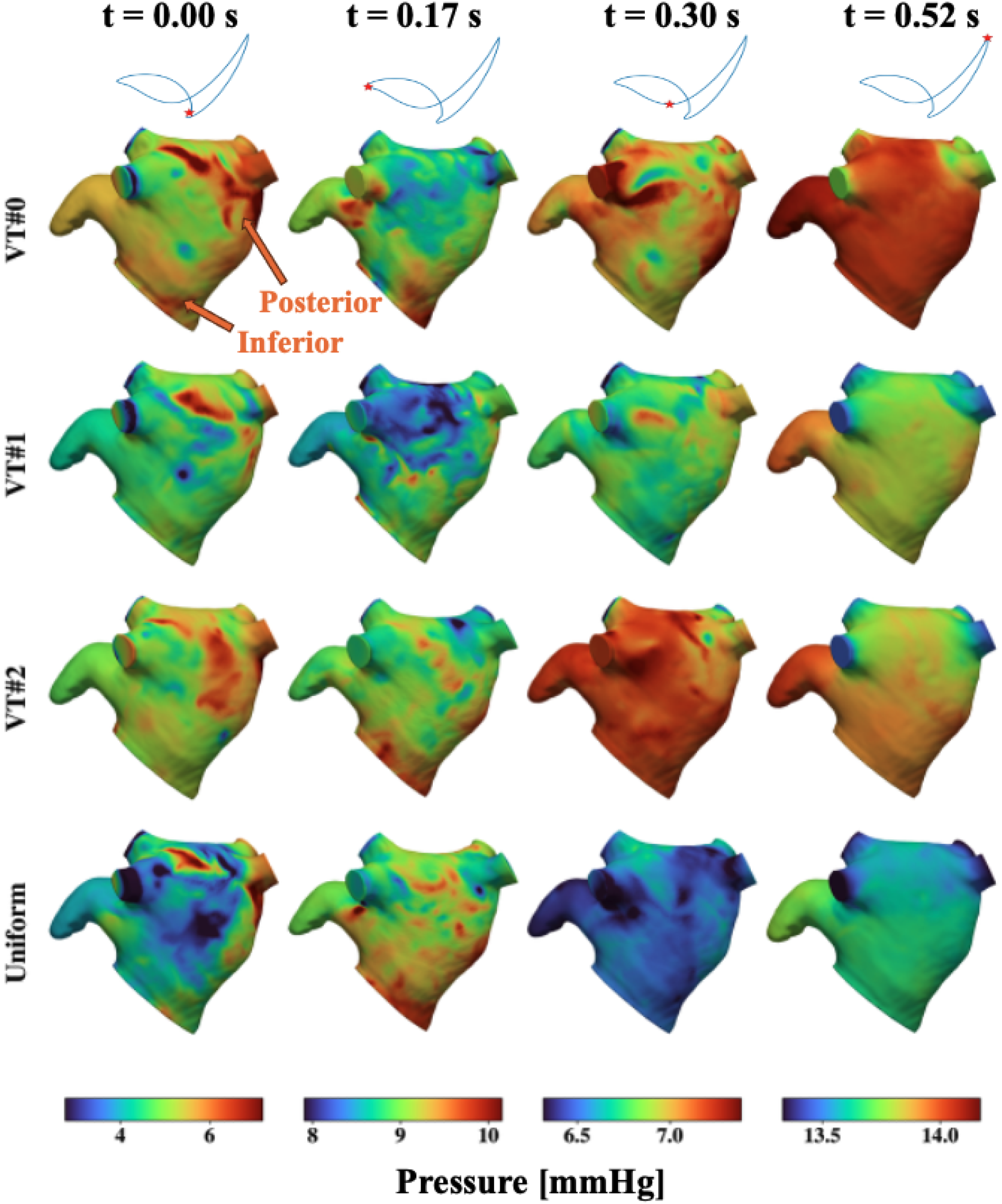
Posterior view of instantaneous intracavitary pressure across the four LA models at four cardiac-cycle phases.

**Fig. A11:**
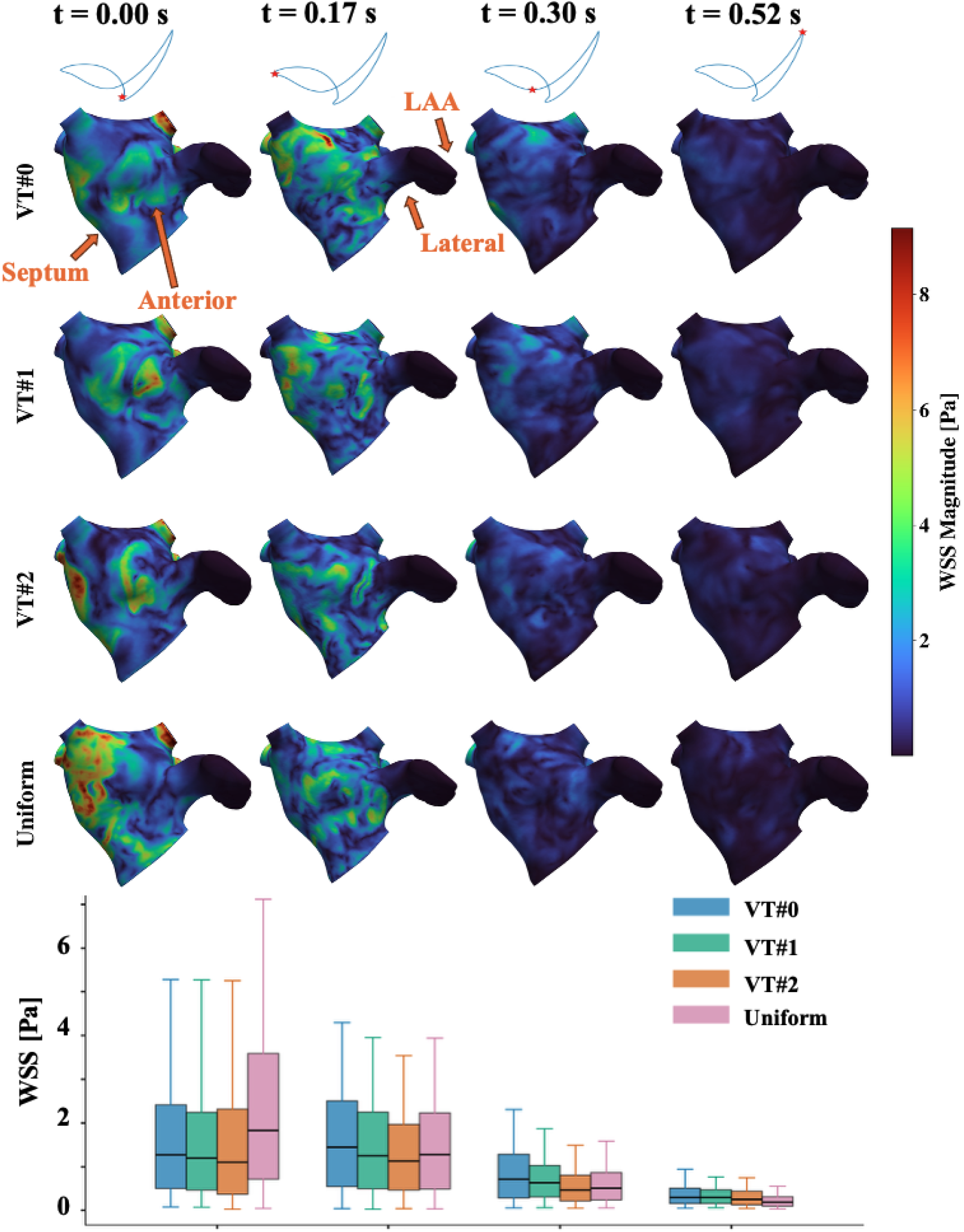
Instantaneous wall shear stress magnitude across the four LA models at four cardiac-cycle phases. Top: posterior surface maps. Bottom: grouped boxplot of displacement of the LA endocardium surface at each phase, with whiskers spanning the 5th–95th percentile.

**Fig. A12:**
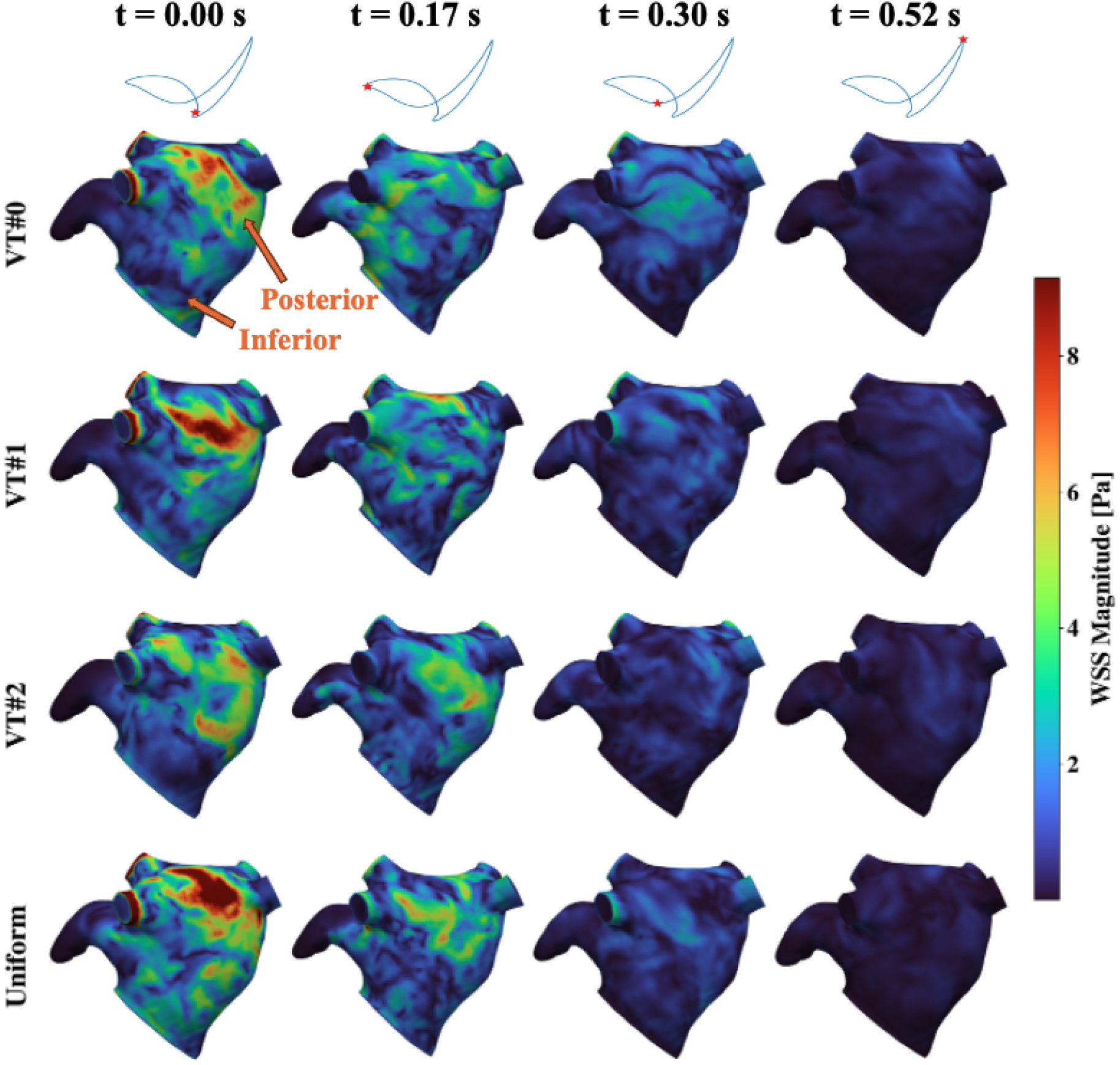
Posterior view of instantaneous wall shear stress magnitude across the four LA models at four cardiac-cycle phases.

**Fig. A13:**
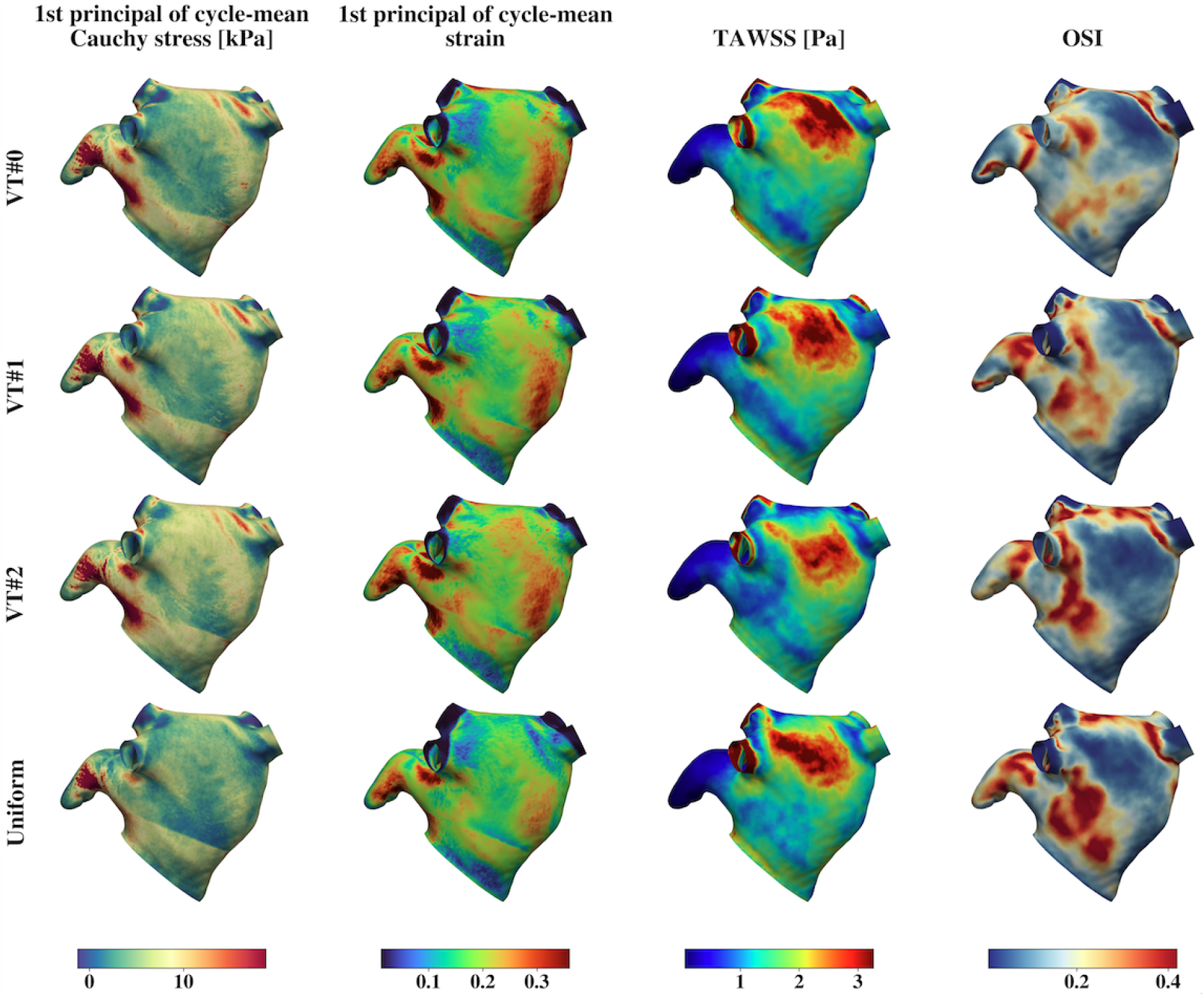
Comparison of time-averaged mechanics and hemodynamics metrics between LAWT variants along the posterior view. The same quantities are compared along the anterior view in Fig. 4. (column-wise) 1st principal Cauchy stress, 1st principal strain, TAWSS, and OSI. (row-wise) LAWT variants including VT#0 (baseline), VT#1, VT#2, and Uniform. LAWT: left atrial wall thickness; TAWSS: time-averaged wall shear stress; OSI: oscillatory shear index.

## References

[1] Stefanadis, C., Dernellis, J., Toutouzas, P.: A clinical appraisal of left atrial function. European Heart Journal 22, 22–36 (2001) 10.1053/euhj.1999.2581

[2] Blume, G.G., McLeod, C.J., Barnes, M.E., Seward, J.B., Pellikka, P.A., Bastiansen, P.M., Tsang, T.S.M.: Left atrial function: physiology, assessment, and clinical implications. European Journal of Echocardiography 12, 421–430 (2011) 10.1093/ejechocard/jeq175

[3] Pagel, P.S., Kehl, F., Gare, M., Hettrick, D.A., Kersten, J.R., Warltier, D.C.: Mechanical function of the left atrium: New insights based on analysis of pressure–volume relations and doppler echocardiography. Anesthesiology 98, 975–994 (2003) 10.1097/00000542-200304000-00027

[4] Ho, S.Y., Sanchez-Quintana, D., Cabrera, J.A., Anderson, R.H.: Anatomy of the left atrium: implications for radiofrequency ablation of atrial fibrillation. Journal of Cardiovascular Electrophysiology 10, 1525–1533 (1999) 10.1111/j.1540-8167.1999.tb00211.x

[5] Ho, S.Y., Cabrera, J.A., Sanchez-Quintana, D.: Left atrial anatomy revisited. Circulation: Arrhythmia and Electrophysiology 5, 220–228 (2012) 10.1161/CIRCEP.111.962720

[6] Al-Saady, N.M., Obel, O.A., Camm, A.J.: Left atrial appendage: structure, function, and role in thromboembolism. Heart 82, 547–554 (1999) 10.1136/hrt.82.5.547

[7] Cheng, S., He, J., Han, Y., Han, S., Li, P., Liao, H., Guo, J.: Global burden of atrial fibrillation/atrial flutter and its attributable risk factors from 1990 to 2021. Europace 26(7), 195 (2024) 10.1093/europace/euae195

[8] Kornej, J., Börschel, C.S., Benjamin, E.J., Schnabel, R.B.: Epidemiology of Atrial Fibrillation in the 21st Century: Novel Methods and New Insights. Lippincott Williams and Wilkins (2020). 10.1161/CIRCRESAHA.120.316340

[9] Vinciguerra, M., Dobrev, D., Nattel, S.: Atrial fibrillation: pathophysiology, genetic and epigenetic mechanisms. Elsevier Ltd (2024). 10.1016/j.lanepe.2023.100785

[10] Choi, S.E., Sagris, D., Hill, A., Lip, G.Y.H., Abdul-Rahim, A.H.: Atrial fibrillation and stroke. Expert Review of Cardiovascular Therapy 21, 35–56 (2023) 10.1080/14779072.2023.2160319

[11] Watson, T., Shantsila, E., Lip, G.Y.H.: Mechanisms of thrombogenesis in atrial fibrillation: Virchow’s triad revisited. The Lancet 373, 155–166 (2009) 10.1016/S0140-6736(09)60040-4

[12] Freedman, B., Potpara, T.S., Lip, G.Y.H.: Stroke prevention in atrial fibrillation. The Lancet 388, 806–817 (2016) 10.1016/S0140-6736(16)31257-0

[13] Boyle, P.M., Alamo, J.C.D., Akoum, N.: Fibrosis, atrial fibrillation and stroke: clinical updates and emerging mechanistic models. Heart 107, 99–105 (2021) 10.1136/heartjnl-2020-316838

[14] Qureshi, A., Balmus, M., Lip, G., Williams, S., Nordsletten, D.A., Aslanidi, O., Vecchi, A.D.: Mechanistic modelling of virchow’s triad to assess thrombogenicity and stroke risk in atrial fibrillation patients. European Heart Journal - Digital Health 3, 278–288 (2022) 10.1093/ehjdh/ztac076

[15] Zingaro, A., Ahmad, Z., Kholmovski, E., Sakata, K., Dede, L., Morris, A.K., Quarteroni, A., Trayanova, N.A.: A comprehensive stroke risk assessment by combining atrial computational fluid dynamics simulations and functional patient data. Scientific Reports 14, 9515 (2024) 10.1038/s41598-024-59997-2

[16] Telle, Å., Bargellini, C., Chahine, Y., Alamo, J.C., Akoum, N., Boyle, P.M.: Personalized biomechanical insights in atrial fibrillation: Opportunities and challenges. Expert Review of Cardiovascular Therapy 21, 817–837 (2023) 10.1080/14779072.2023.2273896

[17] Kjeldsberg, H.A., Albors, C., Mill, J., Medel, D.V., Camara, O., Sundnes, J., Valen-Sendstad, K.: Impact of left atrial wall motion assumptions in fluid simulations on proposed predictors of thrombus formation. International Journal for Numerical Methods in Biomedical Engineering 40, 3825 (2024) 10.1002/cnm.3825

[18] Brown, A.L., Shi, L., Salvador, M., Kong, F., Ennis, D.B., Chen, I., Vedula, V., Marsden, A.L.: Personalized biventricular mechanics and sensitivity to model morphology (2025). https://doi.org/10.64898/2025.12.11.693778. 10.64898/2025.12.11.693778

[19] Vecchi, A., Camara, O., Cavarra, R., Alamo, J.C., El-Bouri, W., Ferro, A., Lu, H.H.-S., Melidoro, P., Ogbomo-Harmitt, S., Olier, I., Ortega-Martorell, S., Patell, R., Vergara, C., Volpert, V., Lip, G.Y.H., Aslanidi, O.: Digital twins for predictive modelling of thrombosis and stroke risk: Current approaches and future directions. Thrombosis and Haemostasis (2026) 10.1055/a-2761-5903

[20] Telle, Å., Maleckar, M.M., Wall, S.T., Powers, J.D., Augustin, C.M., Sundnes, J., Boyle, P.M.: Mechanical modeling of cardiac fibrosis with explicit spatial representation of cellular structure and collagen alignment. Journal of Biomechanical Engineering 148, 021007 (2026) 10.1115/1.4070346

[21] Shi, L., Chen, I.Y., Takayama, H., Vedula, V.: An optimization framework to personalize passive cardiac mechanics. Computer Methods in Applied Mechanics and Engineering 432 (2024) 10.1016/j.cma.2024.117401

[22] Shi, L., Gan, B., Chen, I.Y., Vedula, V.: Personalized multiscale modeling of left atrial mechanics and blood flow. Computer Methods in Applied Mechanics and Engineering 448 (2026) 10.1016/j.cma.2025.118412

[23] Vedula, V., George, R., Younes, L., Mittal, R.: Hemodynamics in the left atrium and its effect on ventricular flow patterns. Journal of Biomechanical Engineering 137, 111003 (2015) 10.1115/1.4031487

[24] Otani, T., Al-Issa, A., Pourmorteza, A., McVeigh, E.R., Wada, S., Ashikaga, H.: A computational framework for personalized blood flow analysis in the human left atrium. Annals of Biomedical Engineering 44, 3284–3294 (2016) 10.1007/s10439-016-1590-x

[25] Dillon-Murphy, D., Marlevi, D., Ruijsink, B., Qureshi, A., Chubb, H., Kerfoot, E., O’Neill, M., Nordsletten, D., Aslanidi, O., Vecchi, A.D.: Modeling left atrial flow, energy, blood heating distribution in response to catheter ablation therapy. Frontiers in Physiology 9, 1757 (2018) 10.3389/fphys.2018.01757

[26] Masci, A., Barone, L., Dede, L., Fedele, M., Tomasi, C., Quarteroni, A., Corsi, C.: The impact of left atrium appendage morphology on stroke risk assessment in atrial fibrillation: a computational fluid dynamics study. Frontiers in Physiology 9, 1938 (2019) 10.3389/fphys.2018.01938

[27] Masci, A., Alessandrini, M., Forti, D., Menghini, F., Dedè, L., Tomasi, C., Quarteroni, A., Corsi, C.: A proof of concept for computational fluid dynamic analysis of the left atrium in atrial fibrillation on a patient-specific basis. Journal of Biomechanical Engineering 142, 011002 (2020) 10.1115/1.4044583

[28] Garcia-Villalba, M., Rossini, L., Gonzalo, A., Vigneault, D., Martinez-Legazpi, P., Duran, E., Flores, O., Bermejo, J., McVeigh, E., Kahn, A.M., Vecchi, A.D.: Demonstration of patient-specific simulations to assess left atrial appendage thrombogenesis risk. Frontiers in Physiology 12, 596596 (2021) 10.3389/fphys.2021.596596

[29] Paliwal, N., Ali, R.L., Salvador, M., O’Hara, R., Yu, R., Daimee, U.A., Akhtar, T., Pandey, P., Spragg, D.D., Calkins, H., Trayanova, N.A.: Presence of left atrial fibrosis may contribute to aberrant hemodynamics and increased risk of stroke in atrial fibrillation patients. Frontiers in Physiology 12, 657452 (2021) 10.3389/fphys.2021.657452

[30] Qureshi, A., Darwish, O., Dillon-Murphy, D., Chubb, H., Williams, S., Nechipurenko, D., Ataullakhanov, F., Nordsletten, D., Aslanidi, O., Vecchi, A.D.: Modelling left atrial flow and blood coagulation for risk of thrombus formation in atrial fibrillation. In: 2020 Computing in Cardiology, pp. 1–4. IEEE, Piscataway, NJ, USA (2020). 10.22489/CinC.2020.219

[31] Gonzalo, A., Augustin, C.M., Bifulco, S.F., Telle, Å., Chahine, Y., Kassar, A., Guerrero-Hurtado, M., Durán, E., Martínez-Legazpi, P., Flores, O., Bermejo, J., Plank, G., Akoum, N., Boyle, P.M., Alamo, J.C.: Multiphysics simulations reveal haemodynamic impacts of patient-derived fibrosis-related changes in left atrial tissue mechanics. The Journal of Physiology 602, 6789–6812 (2024) 10.1113/JP287011

[32] Rodríguez-Aparicio, S., Ferrera, C., Fuentes-Cañamero, M.E., García, J.G., Dueñas-Pamplona, J.: Morphing the left atrium geometry: The role of the pulmonary veins on flow patterns and thrombus formation. Computers in Biology and Medicine 186, 109612 (2025) 10.1016/j.compbiomed.2024.109612

[33] Qureshi, A., Melidoro, P., Balmus, M., Lip, G.Y.H., Nordsletten, D.A., Williams, S.E., Aslanidi, O., Vecchi, A.: MRI-based modelling of left atrial flow and coagulation to predict risk of thrombogenesis in atrial fibrillation. Medical Image Analysis 101, 103475 (2025) 10.1016/j.media.2025.103475

[34] Feng, L., Gao, H., Griffith, B., Niederer, S., Luo, X.: Analysis of a coupled fluid-structure interaction model of the left atrium and mitral valve. International Journal for Numerical Methods in Biomedical Engineering 35, 3254 (2019) 10.1002/cnm.3254

[35] Baptiste, T.M.G., Rodero, C., Sillett, C.P., Strocchi, M., Lanyon, C.W., Augustin, C.M., Lee, A.W.C., Solís-Lemus, J.A., Roney, C.H., Ennis, D.B., Rajani, R., Rinaldi, C.A., Plank, G., Wilkinson, R.D., Williams, S.E., Niederer, S.A.: Regional heterogeneity in left atrial stiffness impacts passive deformation in a cohort of patient-specific models. PLoS Computational Biology 21, 1013656 (2025) 10.1371/journal.pcbi.1013656

[36] Telle, Å., Kassar, A., Chamoun, N., Haykal, R., Gonzalo, A., Hensley, T., Chahine, Y., Flores, O., Álamo, J.C., Akoum, N., Augustin, C.M., Boyle, P.M.: Systematic computational assessment of atrial function impairment due to fibrotic remodeling in electromechanical properties. PLOS Computational Biology 21, 1013265 (2025) 10.1371/journal.pcbi.1013265

[37] Varela, M., Morgan, R., Theron, A., DIllon-Murphy, D., Chubb, H., Whitaker, J., Henningsson, M., Aljabar, P., Schaeffter, T., Kolbitsch, C., Aslanidi, O.V.: Novel mri technique enables non-invasive measurement of atrial wall thickness. IEEE Transactions on Medical Imaging 36, 1607–1614 (2017) 10.1109/TMI.2017.2671839

[38] Zhao, J., Hansen, B.J., Csepe, T.A., Lim, P., Wang, Y., Williams, M., Mohler, P.J., Janssen, P.M.L., Weiss, R., Hummel, J.D., Fedorov, V.V.: Integration of High-Resolution Optical Mapping and 3-Dimensional Micro-Computed Tomographic Imaging to Resolve the Structural Basis of Atrial Conduction in the Human Heart. Lippincott Williams and Wilkins (2015). 10.1161/CIRCEP.115.003064

[39] Beinart, R., Abbara, S., Blum, A., Ferencik, M., Heist, E.K., Ruskin, J., Mansour, M.: Left atrial wall thickness variability measured by ct scans in patients undergoing pulmonary vein isolation. Journal of Cardiovascular Electrophysiology 22, 1232–1236 (2011) 10.1111/j.1540-8167.2011.02100.x

[40] Bishop, M., Rajani, R., Plank, G., Gaddum, N., Carr-White, G., Wright, M., O’Neill, M., Niederer, S.: Three-dimensional atrial wall thickness maps to inform catheter ablation procedures for atrial fibrillation. Europace 18, 376–383 (2016) 10.1093/europace/euv073

[41] Takahashi, K., Okumura, Y., Watanabe, I., Nagashima, K., Sonoda, K., Sasaki, N., Kogawa, R., Iso, K., Ohkubo, K., Nakai, T., Hirayama, A.: Relation between left atrial wall thickness in patients with atrial fibrillation and intracardiac electrogram characteristics and atp-provoked dormant pulmonary vein conduction. Journal of Cardiovascular Electrophysiology 26, 597–605 (2015) 10.1111/jce.12660

[42] Karim, R., Blake, L.-E., Inoue, J., Tao, Q., Jia, S., Housden, R.J., Bhagirath, P., Duval, J.-L., Varela, M., Behar, J.M., Cadour, L., Geest, R.J., Cochet, H., Drangova, M., Sermesant, M., Razavi, R., Aslanidi, O., Rajani, R., Rhode, K.: Algorithms for left atrial wall segmentation and thickness – evaluation on an open-source ct and mri image database. Medical Image Analysis 50, 36–53 (2018) 10.1016/j.media.2018.08.004

[43] Hoffmeister, P.S., Chaudhry, G.M., Mendel, J., Almasry, I., Tahir, S., Marchese, T., Haffajee, C.I., Orlov, M.V.: Evaluation of left atrial and posterior mediastinal anatomy by multidetector helical computed tomography imaging: Relevance to ablation. Journal of Interventional Cardiac Electrophysiology 18, 217–223 (2007) 10.1007/s10840-007-9096-y

[44] Hall, B., Jeevanantham, V., Simon, R., Filippone, J., Vorobiof, G., Daubert, J.: Variation in left atrial transmural wall thickness at sites commonly targeted for ablation of atrial fibrillation. Journal of Interventional Cardiac Electrophysiology 17, 127–132 (2007) 10.1007/s10840-006-9052-2

[45] Martino, E.S.D., Bellini, C., Schwartzman, D.S.: In vivo porcine left atrial wall stress: computational model. Journal of Biomechanics 44, 2589–2594 (2011) 10.1016/j.jbiomech.2011.07.021

[46] Hunter, R.J., Liu, Y., Lu, Y., Wang, W., Schilling, R.J.: Left atrial wall stress distribution and its relationship to electrophysiologic remodeling in persistent atrial fibrillation. Circulation: Arrhythmia and Electrophysiology 5, 351–360 (2012) 10.1161/CIRCEP.111.965541

[47] Augustin, C.M., Fastl, T.E., Neic, A., Bellini, C., Whitaker, J., Rajani, R., O’Neill, M.D., Bishop, M.J., Plank, G., Niederer, S.A.: The impact of wall thickness and curvature on wall stress in patient-specific electromechanical models of the left atrium. Biomechanics and Modeling in Mechanobiology 19, 1015–1034 (2020) 10.1007/s10237-019-01268-5

[48] Masci, A., Alessandrini, M., Forti, D., Menghini, F., Dedè, L., Tomasi, C., Quarteroni, A., Corsi, C.: A patient-specific computational fluid dynamics model of the left atrium in atrial fibrillation: development and initial evaluation. In: Functional Imaging and Modeling of the Heart, pp. 392–400. Springer, Cham (2017). 10.1007/978-3-319-59448-4_37

[49] Fanni, B.M., Capellini, K., Di Leonardo, M., Clemente, A., Cerone, E., Berti, S., Celi, S.: Correlation between LAA morphological features and computational fluid dynamics analysis for non-valvular atrial fibrillation patients. Applied Sciences 10, 1448 (2020) 10.3390/app10041448

[50] Kjeldsberg, H.A., Harrison, J., Sermesant, M., Schnabel, R.B., Sundnes, J., Valen-Sendstad, K.: Beyond CHA2DS2-VASc: Hemodynamic and morphologic discriminants for thrombus formation and stroke in atrial fibrillation patients. Annals of Biomedical Engineering (2026) 10.1007/s10439-026-04039-3

[51] Chen, I.Y., Vedula, V., Malik, S.B., Liang, T., Chang, A.Y., Chung, K.S., Sayed, N., Tsao, P.S., Giacomini, J.C., Marsden, A.L., et al.: Preoperative computed tomography angiography reveals leaflet-specific calcification and excursion patterns in aortic stenosis. Circulation: Cardiovascular Imaging 14(12), 1122–1132 (2021)

[52] Scientific Computing and Imaging Institute: Seg3D: volumetric image segmentation and visual-ization. http://www.seg3d.org. Seg3D: volumetric image segmentation and visualization. Last accessed: 2025-08-12 (2025)

[53] Starreveld, R., Does, L.J.M.E., Groot, N.M.S.: Anatomical hotspots of fractionated electrograms in the left and right atrium: do they exist? Europace 21, 60–72 (2019) 10.1093/europace/euy059

[54] Roney, C.H., Pashaei, A., Meo, M., Dubois, R., Boyle, P.M., Trayanova, N.A., Cochet, H., Niederer, S.A., Vigmond, E.J.: Universal atrial coordinates applied to visualisation, registration and construction of patient specific meshes. Medical Image Analysis 55, 65–75 (2019) 10.1016/j.media.2019.04.004

[55] Piersanti, R., Africa, P.C., Fedele, M., Vergara, C., Dedè, L., Corno, A.F., Quarteroni, A.: Modeling cardiac muscle fibers in ventricular and atrial electrophysiology simulations. Computer Methods in Applied Mechanics and Engineering 373, 113468 (2021) 10.1016/j.cma.2020.113468

[56] Fastl, T.E., Tobon-Gomez, C., Crozier, A., Whitaker, J., Rajani, R., McCarthy, K.P., Sanchez-Quintana, D., Ho, S.Y., O’Neill, M.D., Plank, G., Bishop, M.J., Niederer, S.A.: Personalized computational modeling of left atrial geometry and transmural myofiber architecture. Medical Image Analysis 47, 180–190 (2018) 10.1016/j.media.2018.04.001

[57] Pashakhanloo, F., Herzka, D.A., Ashikaga, H., Mori, S., Gai, N., Bluemke, D.A., Trayanova, N.A., McVeigh, E.R.: Myofiber architecture of the human atria as revealed by submillimeter diffusion tensor imaging. Circulation: Arrhythmia and Electrophysiology 9, 004133 (2016) 10.1161/CIRCEP.115.004133

[58] Roney, C.H., Bendikas, R., Pashakhanloo, F., Corrado, C., Vigmond, E.J., McVeigh, E.R., Trayanova, N.A., Niederer, S.A.: Constructing a human atrial fibre atlas. Annals of Biomedical Engineering 49, 233–250 (2021) 10.1007/s10439-020-02641-0

[59] Brown, A.L., Salvador, M., Shi, L., Pfaller, M.R., Hu, Z., Harold, K.E., Hsiai, T., Vedula, V., Marsden, A.L.: A modular framework for implicit 3d–0d coupling in cardiac mechanics. Computer Methods in Applied Mechanics and Engineering 421 (2024) 10.1016/j.cma.2024.116764

[60] Stein, K., Tezduyar, T., Benney, R.: Mesh moving techniques for fluid-structure interactions with large displacements. J. Appl. Mech. 70(1), 58–63 (2003) 10.1115/1.1530635

[61] Zhu, C., Vedula, V., Parker, D., Wilson, N., Shadden, S., Marsden, A.: svfsi: a multiphysics package for integrated cardiac modeling. Journal of Open Source Software 7(78), 4118 (2022) 10.21105/joss.04118

[62] Vedula, V., Lee, J., Xu, H., Kuo, C.-C.J., Hsiai, T.K., Marsden, A.L.: A method to quantify mechanobiologic forces during zebrafish cardiac development using 4-d light sheet imaging and computational modeling. PLoS computational biology 13(10), 1005828 (2017) 10.1371/journal.pcbi.1005828

[63] Zhai, H., Chen, Y., Kitada, Y., Takayama, H., Vedula, V.: Hemodynamic Analysis of a Repaired Ascending Aorta with Preserved Aortic Root (2026). https://doi.org/10.64898/2026.01.28.702307. 10.64898/2026.01.28.702307

[64] Bäumler, K., Vedula, V., Sailer, A.M., Seo, J., Chiu, P., Mistelbauer, G., Chan, F.P., Fischbein, M.P., Marsden, A.L., Fleischmann, D.: Fluid–structure interaction simulations of patient-specific aortic dissection. Biomechanics and modeling in mechanobiology 19(5), 1607–1628 (2020) 10.1007/s10237-020-01294-8

[65] Bazzi, M.S., Balouchzadeh, R., Pavey, S.N., Quirk, J.D., Yanagisawa, H., Vedula, V., Wagenseil, J.E., Barocas, V.H.: Experimental and mouse-specific computational models of the fbln4 smko mouse to identify potential biomarkers for ascending thoracic aortic aneurysm. Cardiovascular engineering and technology 13(4), 558–572 (2022) 10.1007/s13239-021-00600-4

[66] Wang, H., Balzani, D., Vedula, V., Uhlmann, K., Varnik, F.: On the potential self-amplification of aneurysms due to tissue degradation and blood flow revealed from fsi simulations. Frontiers in Physiology 12, 785780 (2021) 10.3389/fphys.2021.785780

[67] Wang, H., Uhlmann, K., Vedula, V., Balzani, D., Varnik, F.: Fluid-structure interaction simulation of tissue degradation and its effects on intra-aneurysm hemodynamics. Biomechanics and Modeling in Mechanobiology 21(2), 671–683 (2022) 10.1007/s10237-022-01556-7

[68] Cliff, N.: Dominance statistics: Ordinal analyses to answer ordinal questions. Psychological Bulletin 114, 494–509 (1993) 10.1037/0033-2909.114.3.494

